# Niche-dependent modular regulation of the stem cell transcriptome separates cell identity and potential

**DOI:** 10.1101/2025.04.10.648217

**Authors:** Amelie Raz, Hafidh Hassan, Yukiko Yamashita

**Affiliations:** Whitehead Institute for Biomedical Research, 455 Main St, Cambridge MA 02142; Department of Biology, Massachusetts Institute of Technology, 31 Ames Street Cambridge MA, 02142; Howard Hughes Medical Institute, 455 Main St, Cambridge MA 02142

**Keywords:** Stem cells, dedifferentiation, germline, niche signaling

## Abstract

Adult stem cells maintain tissue homeostasis, yet are themselves vulnerable to loss. One common mechanism to replace lost stem cells is dedifferentiation, in which progeny revert to stem cell identity. It is a paradox how stem cells and progeny retain the same stem cell potential while exhibiting distinct current identities of self-renewal, differentiation, and dedifferentiation. Here, we show that the Drosophila male germline lineage solves this paradox via two parallel and complementary mechanisms to separate potential and identity. First, differentiating progeny maintain stem cell potency by inheriting perdurant stem cell mRNAs without actively transcribing them. Second, two known niche signals (Bmp and Jak-Stat) activate distinct sets of targets, defining three identities (self-renewal, differentiation, and dedifferentiation) based on the combination of their on/off states. Together, this study reveals how a pool of dedifferentiation-competent progeny is maintained to regenerate stem cells as needed without resulting in stem cell overproduction, and resolves the puzzle of why most stem cell systems require multiple independent niche signals.

**Significance Statement:** Dedifferentiation is a mechanism by which differentiating cells revert back to stem cell identity to compensate for the loss of stem cells, allowing for long-term tissue homeostasis. It remains poorly understood how stem cells and progeny retain the same stem cell potential while exhibiting distinct current identities of self-renewal, differentiation, and dedifferentiation. This study reveals two parallel mechanisms to separate cells’ potential and identity. First, differentiating progeny maintain stem cell potency by inheriting perdurant stem cell mRNAs. Second, two known niche signals (Bmp and Jak-Stat) activate distinct sets of targets, defining three identities (self-renewal, differentiation, and dedifferentiation) based on the combination of their on/off states. This study reveals how a pool of dedifferentiation-competent progeny is maintained to regenerate stem cells.

## Introduction

Adult stem cells maintain tissue homeostasis by continuous production of differentiated cells. However, stem cells can be lost to damage. Thus, mechanisms to replace stem cells are critical for long-term tissue homeostasis. Dedifferentiation serves as one such replacement mechanism, in which the process of differentiation is reversed to generate stem cells from partially or fully differentiated progeny^1^. Dedifferentiation has been observed in stem cell lineages of the intestine^2^, germline^3–5^, hair follicle^6^, liver^7,8^, neuron^9^, muscle^10^, airway epithelium^11^, and others. In a system that uses dedifferentiation, stem cells, differentiating cells, and dedifferentiating cells share the same stemness potential – i.e., the ability to be or become stem cells. However, each of these cells has distinct identities: self-renewing stem cells, differentiating progeny, and dedifferentiating cells. It is critically important to separate a cell’s potential (e.g., ability to dedifferentiate) from current identity (e.g., actual dedifferentiation): the inevitable realization of the potential would lead to dysregulation of tissue homeostasis, including tumorigenesis^12^. How cells of distinct identities maintain the same potential remains poorly understood.

The male gonads of the fruit fly *Drosophila melanogaster* are an ideal model for studies of stem cell regulation. The adult Drosophila male germline is composed of germline stem cells (GSCs) and their differentiating progeny, all of which are present in a spatial-temporal order along the length of the adult testis (**Fig 1A**). At the apical tip of the testis, GSCs are physically attached to a cluster of somatic niche cells called the hub, which instructs GSC self-renewal through the secretion of signaling ligands for Bmp and Jak-Stat pathways^13^. GSCs divide asymmetrically, producing one GSC and one differentiating cell, the latter of which continues to migrate away from the niche while undergoing 4 rounds of mitotic divisions with incomplete cytokinesis. These divisions thus yield cysts of 2, 4, 8, or 16 interconnected spermatogonia (each called 2-cell spermatogonia, 4-cell spermatogonia, etc.) (**Fig 1A**)^14^. GSCs and their progeny of each differentiation stage can be easily identified by their position within the tissue and the connectivity. These cells eventually enter meiotic program as spermatocytes, and complete differentiation to become mature sperm.

**Figure 1.**
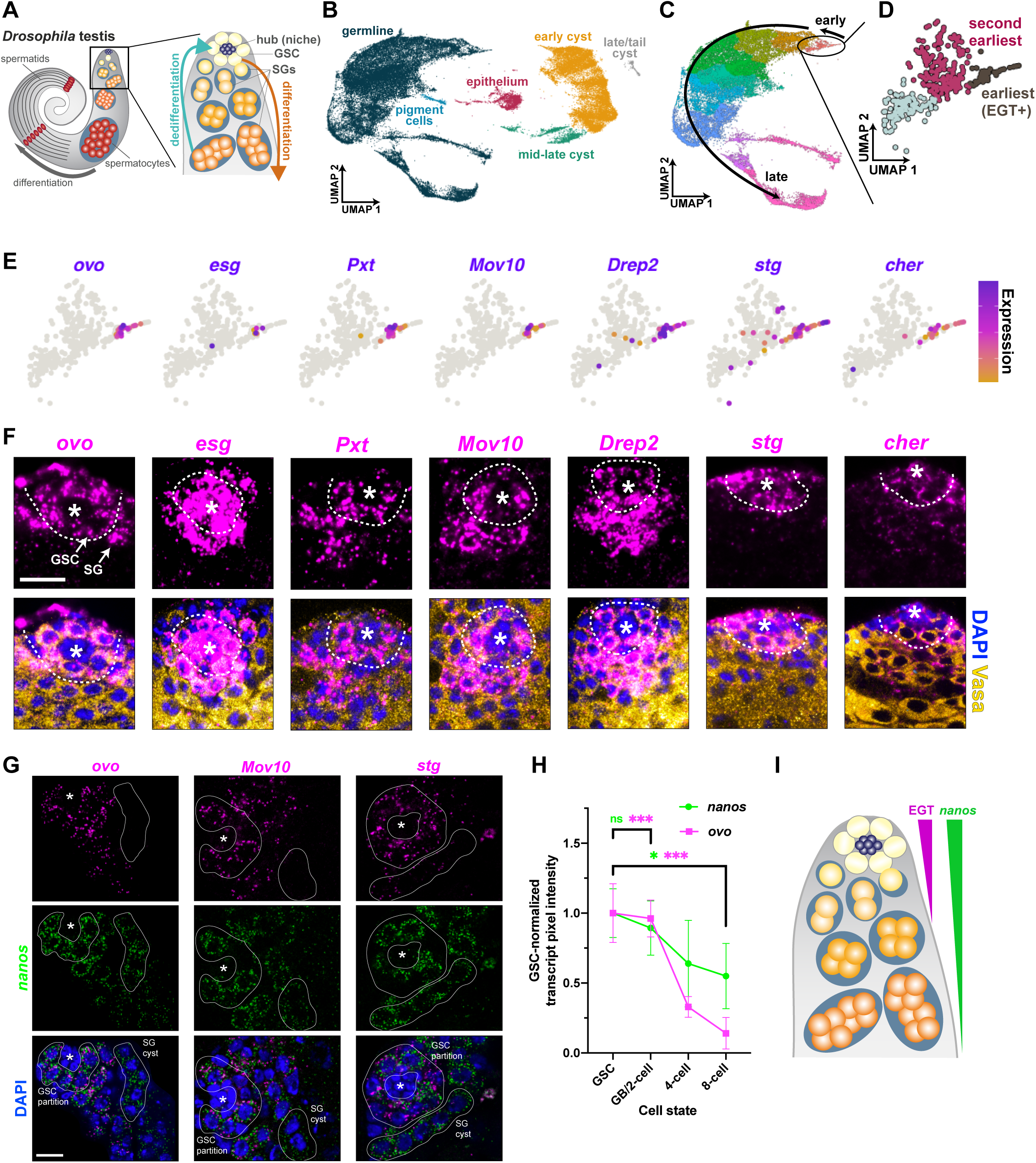
Single-cell analysis reveals an early germline transcriptome (EGTome). **(A)** Schematic of the Drosophila testis. The adult Drosophila male germline is maintained by germline stem cells (GSCs), which can both self-renew and give rise to differentiating spermatogonia. Spermatogonia divide four times while differentiating, mature into pre-meiotic spermatocytes, undergo meiosis as spermatids, and mature as sperm. Spermatogonia can dedifferentiate to GSCs. **(B)** UMAP representation of all singly sequenced cells in the testis. Each dot represents a cell, and the proximity of cells to each other approximates closeness in gene expression space. Color denotes broad cluster annotation. **(C)** UMAP representation of all germline annotated cells in the testis. Arrows indicate differentiation direction. The GSCs and spermatogonia (colored salmon and circled) are collectively known as “gonia”. Color indicates merged and quality-controlled germline cluster annotation. **(D)** UMAP representation of gonia, colored by further subcluster. Single-cell differential expression analysis compared putative spermatogonia (magenta in UMAP plot) to putative GSCs (dark grey-brown in UMAP plot). **(E)** UMAP plots of gonia with representative early germline transcripts (EGTs) featured. Color indicates expression of gene listed, with blue as highest and orange as lowest detectable. Grey cells have no detectable expression. **(F)** RNA FISH of EGTs in 1E. Asterisk marks the hub. All factors are expressed in GSCs (Vasa^+^ cells contacting the hub, within the dashed line) and in at least some spermatogonia (Vasa^+^ cells not contacting the hub, outside the dashed line). Some factors, including *esg, stg,* and *cher* are also expressed in somatic cells. Scale bar, 10 µm. **(G)** Single-molecule FISH of EGTs reveals that all EGTs assayaed are more restricted to GSCs than *nanos* transcript, a common GSC-enriched marker. In all images, both dual-positive GSCs and *nanos*-single positive spermatogonia are highlighted. Asterisk marks the hub. Scale bars, 10 µm. **(H)** Quantification of pixel intensity from **(G)**. Error bars, standard error. *, p<0.01; ***, p<0.0001, unpaired t-test. **(I)** Model of transcript presence in the early germline. High EGT transcript presence is restricted to GSCs through approximately two-cell spermatogonia. *nanos* transcript presence is found in GSCs through 8-cell spermatogonia, in a decreasing gradient.

Notably, GSCs and spermatogonia exhibit quite divergent cell biology, with different cell death, mitotic recombination, replication, cytokinesis, and polarity behaviors^15–21^. These distinct behaviors could be suggestive of a model in which GSCs represent a unique and exclusive cell type. However, such a model is complicated by the fact that spermatogonia frequently and spontaneously undergo dedifferentiation to generate GSCs^22^. Upon experimental depletion of GSCs, spermatogonia were observed to dedifferentiate *en masse* to occupy the empty niche, restoring the GSC pool^23–26^. Early spermatogonia (1-2 cell spermatogonia) comprise the majority of dedifferentiation events, with late spermatogonia (4-16 cell spermatogonia) dedifferentiating less frequently^5^. Dedifferentiation of spermatogonia is required for the maintenance of GSC populations long-term even in unperturbed tissues^5,25^, resulting in >40% of GSCs being generated from dedifferentiation by the end of the animal’s lifespan^27^. GSCs that arise from dedifferentiation return to asymmetric self-renewal divisions and generate progeny with fairly normal kinetics^5,25^. It remains unknown how GSCs and spermatogonia maintain the same potential to be or become stem cells, while maintaining distinct cell identities and exhibiting different behaviors.

Here, we reveal two parallel mechanisms that allow spermatogonia to assume a differentiation identity while maintaining stem cell potential (i.e., the ability to dedifferentiate). First, through analysis of single-cell RNA sequencing, we find that GSCs and early spermatogonia share an indistinguishable transcriptome, which we call the “early germ cell transcriptome” or “EGTome”, suggesting that the separate identity of GSCs and spermatogonia are not defined by the transcriptome. Rather, we find that GSCs are distinguished from spermatogonia by their ability to actively transcribe early germline transcripts (EGTs) that constitute the EGTome, whereas spermatogonia inherit EGTs as perdurant mRNAs. Thus, we propose that spermatogonia preserve stem cell potential through maintenance of EGT mRNAs, while assuming a differentiation identity associated with halting EGT transcription. Second, we find that the identified EGTome is activated in separable subsets by independent activities of the Bmp and Jak-Stat pathways. As a result, the combinatorial activity of the Bmp and Jak-Stat pathways can determine the individual trajectories (self-renewal, differentiation, or dedifferentiation) from a pool of equally-potent cells. Reception of both, neither, or Bmp ligand alone each yields a distinct outcome: self-renewal, differentiation, or dedifferentiation, respectively. We propose that separation of potential and identity via perdurance of GSC-produced transcripts and use of modular transcriptome via multiple signaling pathways serves as a key mechanism to sustain stem cells while avoiding stem cell overproliferation.

## Results

### Identification of GSC-enriched transcripts with single-cell RNA sequencing

Cell identity is generally thought to be represented by a specific transcript profile. However, no GSC-specific transcript has been identified to date^28^. Thus far, single-cell and single-nucleus RNA sequencing have identified a transcriptomic cluster of early germ cells that contains a mixture of GSCs and early spermatogonia^28^. We attempted to distinguish GSCs from spermatogonia by further analyzing transcriptional heterogeneity within the earliest germ cell cluster. In all, we performed single-cell sequencing of over 51,000 quality-controlled cells across six biological replicates of dissociated testes (see methods, **Supp Fig 1**). Clustering^29^ first yielded distinct superclusters representing the primary cell types present in the tissue, including germline and somatic subpopulations (pigment cells, epithelium, and encapsulating cyst cells) (**Fig 1B**, **Supp Fig 1A-B**). More granular clustering of the germline partition, alongside pseudotemporal trajectory inference^29^ (**Supp Fig 1B-C**) (Methods), revealed subclusters with gene expression profiles indicating continuous differentiation from stem cells to mature spermatids (**Fig 1C**, **Supp Fig 1A-E**). Cells at the root of the trajectory (circled and salmon-colored in **Fig 1C**, dark blue in **Supp Fig 1C**) represent germline stem cells and their recent progeny. By extracting these cells and performing a final round of subclustering, we identified the smallest clusters that could be robustly computationally identified (see **Methods**). To determine potential heterogeneity in this cluster representing a distinct transcriptional program for GSCs, we compared the very earliest sub-cluster (**Fig 1D**, brown) with the second-earliest cluster (**Fig 1D**, magenta), using single-cell differential expression (SCDE) analysis optimized for low cell counts^30^. Through this strategy, we identified many transcripts with expression restricted to the earliest germ cells in our dataset (**Fig 1E, Table S1**). These transcripts, referred to collectively as “early germline transcripts” (EGTs), included factors that had not been previously described in the male germline, and several of which encode for transcription factors or RNA binding proteins that regulate cell identity in other contexts^31–33^.

To determine if the computationally-defined ‘earliest cluster’ represented GSCs, we used RNA fluorescent *in situ* hybridization (FISH) to examine the expression patterns of several EGTs (**Table 1**) (*ovo*, *stg*, *cher, esg*, *Mov10*, *Pxt*, and *Drep2*)^31,33–38^. Of the top 30 cluster-enriched transcripts (sorted by conservative estimate, see **Methods**), these seven transcripts were most easily observable via FISH. All of these transcripts were expressed in early germ cells including GSCs and 2-cell spermatogonia, but diminished in 4- and 8-cell spermatogonia (**Fig 1F-H**). This is the most restricted transcriptome reported to date for GSCs and their immediate progeny, compared to previously identified ‘GSC-enriched factors’, such as *nanos,* that exhibit mRNA distribution through 8-cell spermatogonia^39^ (**Fig 1G-I**). Notably, however, no EGTs were restricted to GSCs alone, suggesting that GSCs may not have a unique transcriptome, and instead share their transcriptional profile with early (2-cell) spermatogonia.

### Active EGT transcription distinguishes GSCs from early spermatogonia

The above results revealed that GSCs and their early progeny (through 2-cell spermatogonia) likely share the same early germ cell transcriptome (EGTome), and these two cell types may not be distinguishable via transcript presence. However, it is well-established that the transcriptional activity of the Bmp and Jak-Stat pathways (**Fig 2A**) is restricted to GSCs and daughter cells that have not yet separated from the GSC mother^40–42^ (as confirmed in **Fig 2B-C**): phosphorylated, active form of Mad (pMad) downstream of Bmp, as well as Stat downstream of Jak-Stat, are only detectable in GSCs and their connected daughter cells. In the absence of ligand reception, Mad and Stat are not localized to the nucleus and are therefore not active as transcription factors; conversely, upon ligand reception, Mad (now phosphorylated) and Stat localize to the nucleus and drive gene expression^43,44^. If Mad and Stat are responsible for the expression of EGTs, EGTs’ active transcription may be limited to GSCs, even if mRNAs are inherited and maintained in spermatogonia.

**Figure 2.**
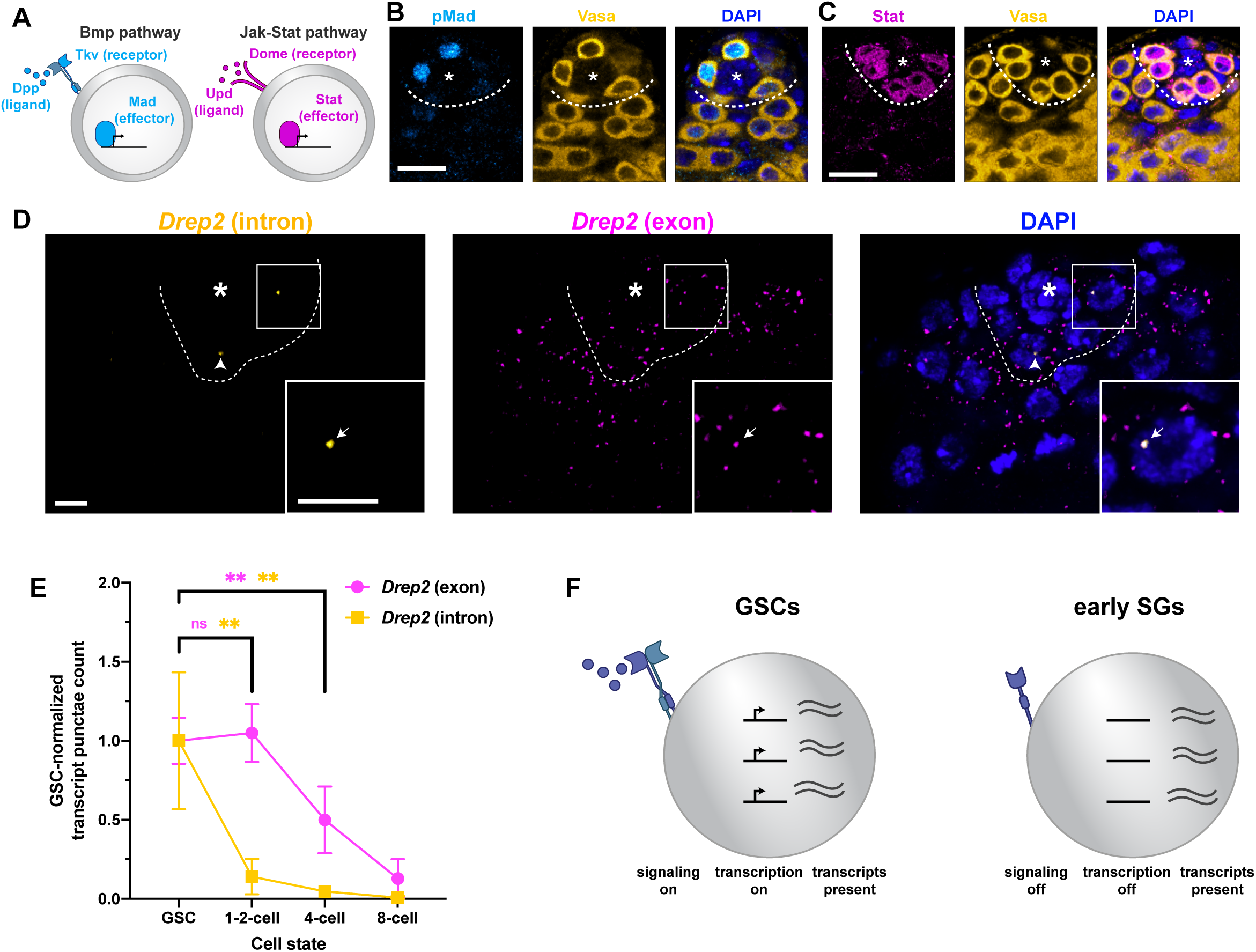
GSCs and spermatognia differ on the basis of active transcriptional state. **(A)** Model of the Bmp and Jak-Stat pathways in Drosophila. In the Bmp pathway, Dpp (ligand) binds to the receptor Tkv, triggering the nuclear localization of the transcription factor Mad. In the Jak-Stat pathway, Upd (ligand) binds to the receptor Dome, triggering nuclear localization of the transcription factor Stat. **(B)** GSCs, but not spermatogonia, are positive for the Bmp transcriptional effector Mad. Scale bar, 10 µm. **(C)** GSCs, but not spermatogonia, are positive for the Jak-Stat transcriptional effector Stat. Scale bar, 10 µm. **(D)** Nascent transcription can be identified by intronic signal. Intronic signal of EGT *Drep2* is localized to the nucleus of GSCs, whereas exonic signal is present in the nucleus and cytoplasm of both GSCs and spermatogonia. Intronic signal colocalizes with exonic signal. Scale bars, 5 µm. **(E)** Quantification of the data in C. Error bars represent standard error. **, p<0.001, unpaired t-test. **(F)** Model of GSC vs. spermatogonia difference. GSCs receive signaling information from the hub, promoting transcription of EGTs. Spermatogonia do not receive signaling and thus do not have active transcription, but retain mature mRNAs transcribed in GSCs.

Consistent with this notion, we found that nascent transcripts of EGT *Drep2* was limited to GSCs, suggesting that active EGT transcription may be exclusive to GSCs. Nascent transcripts can be specifically identified by RNA *in situ* hybridization against intronic sequences. Of the EGTs (**Figure 1E**), only *Drep2* had a sufficiently long intron to successfully visualize intronic RNA through single-molecule RNA FISH (**Fig 2D**). Intronic RNA FISH was combined with exonic RNA FISH to compare the presence of nascent transcription vs. mature mRNA. Notably, intronic *Drep2* signal was strongly restricted to the nuclei of GSCs alone, whereas exonic signal was present in the nucleus and cytoplasm of GSCs and early spermatogonia (high expression through 2-cell spermatogonia, and low expression in 4-8 cell spermatogonia) (**Fig 2D-E**). Although other EGTs do not have long enough introns to conduct a similar analysis to detect active transcription, we found that the signal of the exonic probes for two other EGTs, *ovo* and *stg,* can be found in the nucleus specifically in GSCs, but not in spermatogonia, whereas cytoplasmic transcripts were present in both GSCs and spermatogonia (**Supp Fig 2A**). These observations are consistent with a model in which EGTs are actively transcribed in GSCs, while perdurant EGT mRNAs are inherited by early spermatogonia. Similar transcript perdurance from GSCs to spermatogonia has been previously shown for *nanos* mRNA^39^, which, unlike EGTs, perdures up to 8-cell spermatogonia. Taken together, we propose that maintenance of the EGTome without their active transcription may allow early spermatogonia to maintain the potential of dedifferentiation while in the process of committing to differentiation (**Fig 2F**). Such early spermatogonia (1-2 cell spermatogonia) have indeed been shown to be especially dedifferentiation competent relative to later (EGT-negative) spermatogonia^5,45^, suggesting that early spermatogonia may more readily dedifferentiate because of their maintained EGTome.

### Active transcription of EGTs is largely regulated by niche signaling

Although GSC-specific active transcription could only be directly shown for *Drep2* due to the lack of sizeable introns within other EGTs, we hypothesized that transcription of other EGTs may be under the control of GSC-specific Bmp and Jak-Stat signaling. Consistent with this hypothesis, we found that most EGTs examined are upregulated by the activation of Bmp or Jak-Stat pathway (**Fig 3A-D**), suggesting that transcription of EGTs is indeed regulated by the niche signaling. It is well-established that Dpp overexpression results in tumorous overproliferation of spermatogonia-like cells (referred to as Bmp tumors hereafter)^46^, whereas Upd overexpression results in massive overproliferation of GSC-like cells (referred to as Jak-Stat tumors hereafter)^47^. In these tumorous testes, we found that the majority of EGTs are indeed upregulated (**Fig 3A-D**), supporting our hypothesis that active transcription of EGTs is driven by Bmp and/or Jak-Stat signaling and thus typically restricted to GSCs. We specifically examined the *ovo* transcript, which was found to be activated by Jak-Stat pathway (**Figure 3C, 3E**). We found that *ovo* mRNA was present through the two-cell spermatogonia, outlasting active Stat; this further supports the notion that EGTs are maintained in early spermatogonia via perdurance of RNAs rather than active transcription (**Fig 3E**). Two EGTs (*esg* and *cher*) were not upregulated either by Bmp or Jak-Stat activation, and may require both signaling pathways or an alternative mechanism of promoting transcription (**Fig 3D**). Although we could only detect active Stat in GSCs (**Fig 2C**), prior work has suggested Stat signaling may occur in somatic stem cells and influence GSC behavior non-autonomously^48^. Therefore, we promoted GSC-autonomous Stat signaling by driving expression of a constitutively active form of the JAK Kinase (termed Hopscotch). These tumors recapitulated EGT expression found in Upd overexpression tumors, implying that Stat-dependent EGT expression is driven autonomously in GSCs (**Supp Fig 3A**). Taken together, these results suggest that the majority of the EGTome is transcribed in response to GSC-limited niche signaling, and that GSCs and spermatogonia are distinguished by niche-dependent active transcription of the EGTome.

**Figure 3.**
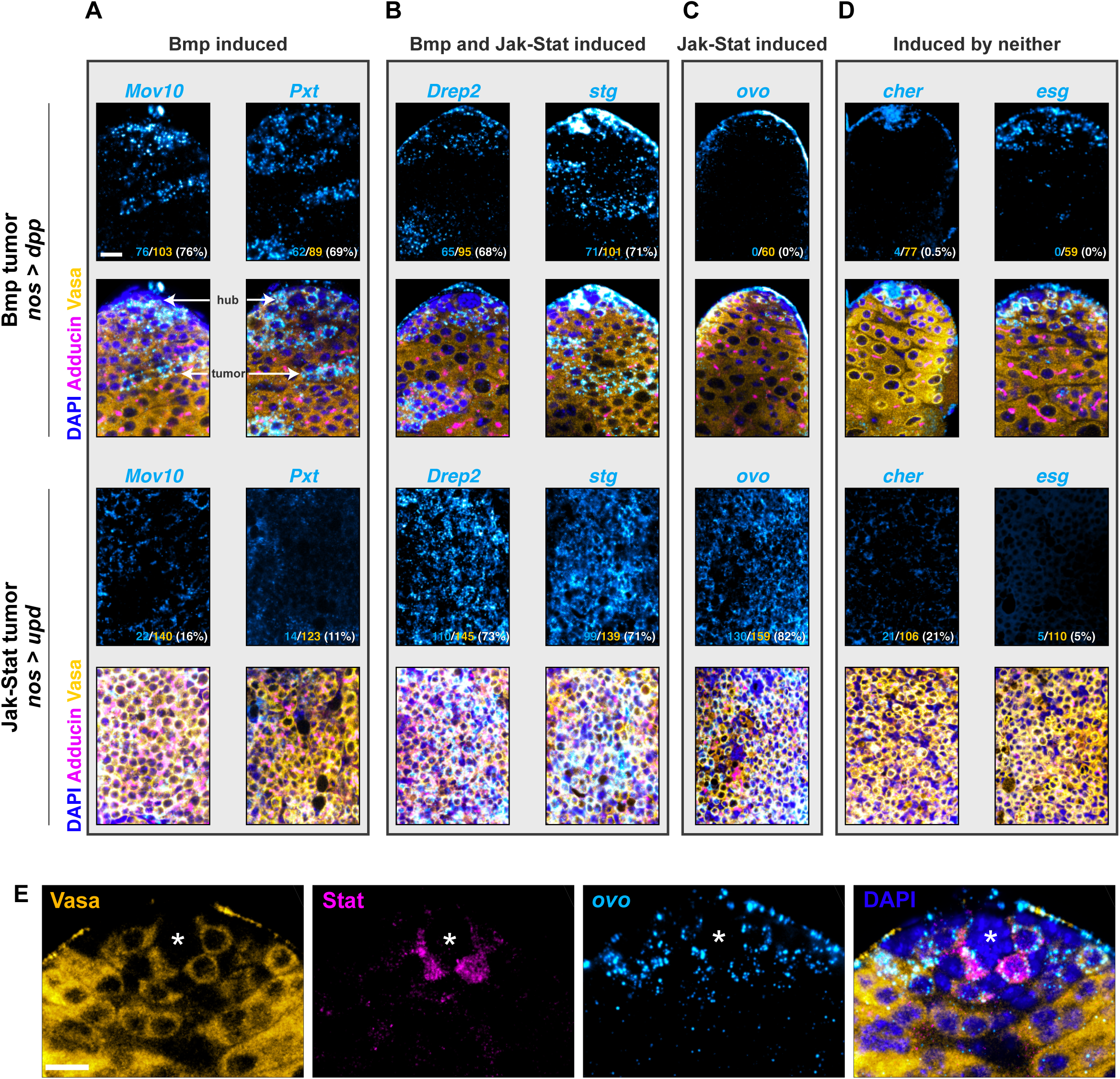
Bmp and Jak-Stat promote separable transcriptional programs. **(A-D)** EGTs have variable expression in conditions of Bmp vs Jak-Stat upregulation. In these testes, hub-proximal germ cells likely represent GSCs receiving both niche ligands from endogenous niche cells, whereas hub-distal tumor cells arise from upregulation of Bmp signaling or Jak-Stat signaling alone, depending on ectopically expressed ligands. **(A)** *Mov10* and *Pxt* are expressed in Bmp tumors, but not Jak-Stat tumors. **(B)** *Drep2* and *stg* are expressed in both Bmp and Jak-Stat tumors. **(C)** *ovo* is expressed in Jak-Stat tumors, but not Bmp tumors. **(D)** *cher* and *esg* are not expressed in either Bmp or Jak-Stat tumors. **(E)** *ovo* transcript presence perdures past Stat transcription factor presence. Stat transcription factor is only found in the nucleus of GSCs, whereas *ovo* transcript, which is sensitive to Stat signaling, is found in early spermatogonia as well. Scale bars, 10 µm.

### Bmp and Jak-Stat signaling act independently to define GSC identity

The above results that showed that EGTs are under the regulation of the niche signaling also surprisingly revealed that Bmp and Jak-Stat tumors differed in their EGT expression profiles. For example, some EGTs (e.g. *Mov10*) were upregulated only in response to Bmp activation, whereas others (e.g. *ovo*) were upregulated only in response to Jak-Stat (**Fig 3A-D, Supp Fig 2B)**. Yet other EGTs (e.g. *stg*) were upregulated in response to either Bmp activation or Jak-Stat activation. These results suggest that the Bmp and Jak-Stat pathways activate overlapping but distinct subsets of the EGTome and operate independently, rather than redundantly.

Supporting the idea that Bmp and Jak-Stat signaling act independently, previous studies have also suggested that each pathway likely regulates distinct aspects of GSC behaviors, although there may be some cross-talk between these two pathways^49^. Specifically, prior work has shown that the tumor phenotypes induced by overexpression of Upd and Dpp differ significantly from each other^40,41,47^. For example, Bmp tumor and Jak-Stat tumor cells demonstrated distinct morphology of the ‘fusome’, the membranous structure that bridges the cytoplasm of differentiating germ cells and has been used to assess the state of germ cell differentiation^20,21^ (**Supp Fig 3B**). Whereas Jak-Stat tumors had ‘spherical fusomes’ (or ‘spectrosome’), Bmp tumors exhibit highly branched fusome morphology spanning large germ cell cysts, indicating that these tumor states are distinct (**Supp Fig 3B**). Mirroring the different states of connectivity, germ cells in Jak-Stat tumors divide asynchronously from their neighbors, whereas germ cells in Bmp tumors share cytoplasm within a cyst and therefore divide synchronously within cysts (**Supp Fig 3C**).

We further confirmed the independence of Bmp and Jak-Stat pathway by demonstrating near-lack of mutual activation by the two pathways. Overexpression of Bmp ligand Dpp caused germ cell tumors with pMad^+^ cells far from the niche (**Figure 4A**), without activating Stat in these tumors (**Fig 4B**); the only Stat+ cells are juxtaposed to the hub cells, likely due to the endogenous niche-secreted Upd ligand. Likewise, overproliferated germ cells caused by Upd overexpression (Jak-Stat tumor) exhibited only a small number of pMad-positive germ cells, mostly around hub-like structures (**Figure 4C**). In these same Jak-Stat tumors, almost all germ cells were positive for Stat (**Fig 4D**). These results suggest that the Bmp and Jak-Stat pathways indeed act independently. Moreover, we found that overexpression of the Jak-Stat ligand Upd does not require Bmp activity to form a tumor. In simultaneous RNAi of the Bmp receptor Thickveins (Tkv) (*tkv^RNAi^*) and overexpression of Jak-Stat ligand Upd, tumors comprised of GSC-like cells formed, just as they do in overexpression of Upd alone (**Supp Fig 4A-B**). These *tkv^RNAi^*, Upd-overexpression tumor did not display nuclear pMad in any germ cells, confirming effective knockdown of *tkv* (**Supp Fig 4B**). Thus, germ cell reception of Bmp signaling is not required for the formation of the Jak-Stat tumor.

**Figure 4.**
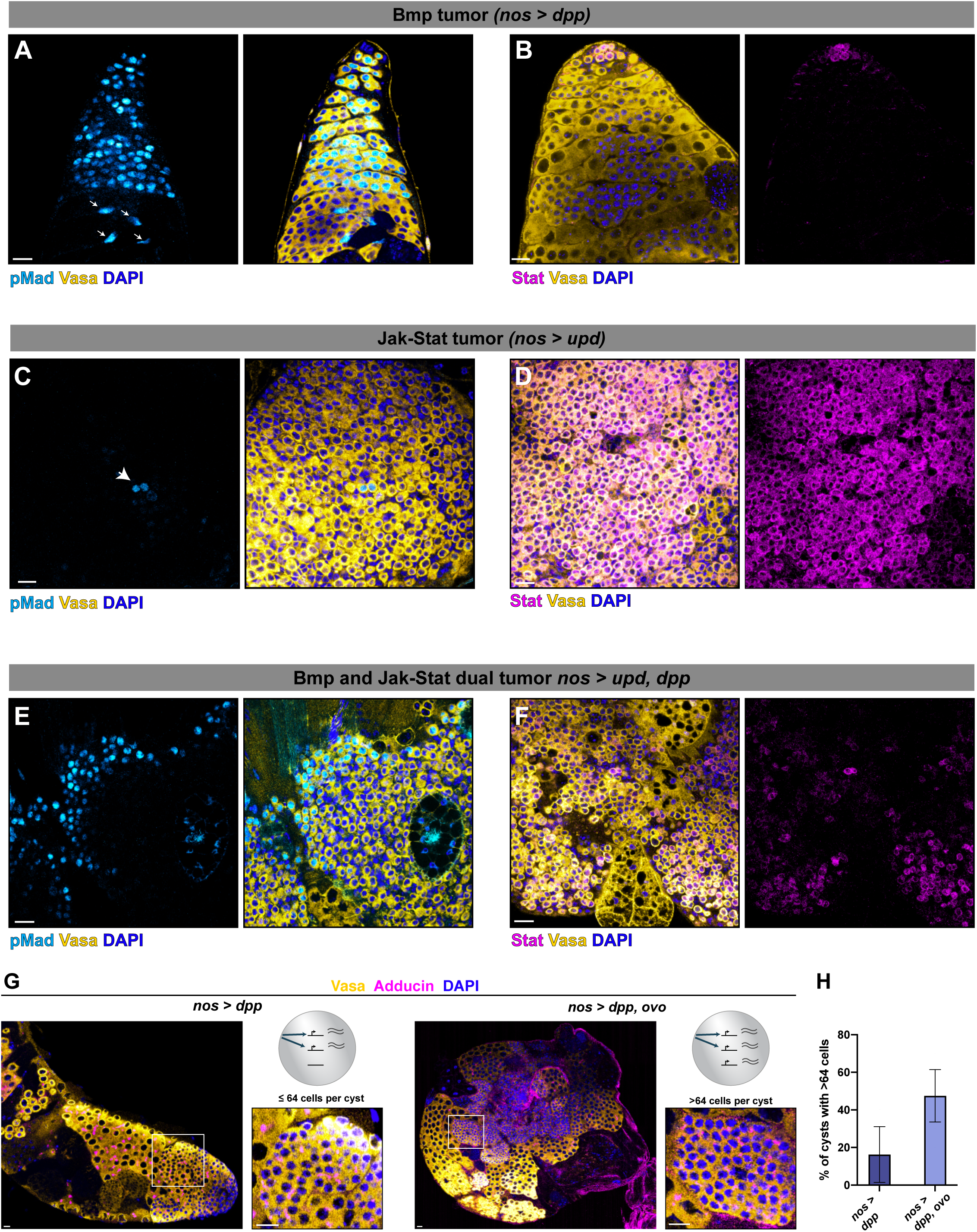
Bmp and Jak-Stat signaling act independently and non-redundantly. **(A)** Bmp tumor cells distal to the hub are pMad^+^. Additionally, some somatic cyst cells distal to the hub are pMad^+^ (arrows). **(B)** Bmp tumor cells distal to the hub are Stat **^-^**. Only hub-proximal cells are Stat^+^. **(C)** Most cells in Jak-Stat tumors are pMad**^-^**. pMad^+^ cells in Jak-Stat tumors are located around hub-like structures (arrow). **(D)** Jak-Stat tumor cells largely are Stat^+^. **(E-F)** Tumors arising from simultaneous overexpression of Upd and Dpp contain many dying or grossly abnormal cells. However, many germ cells are pMad^+^ **(E)** and/or Stat^+^ **(F)**. **(G)** Bmp tumors display tumor cysts distal to the hub with up to 64 cells per cyst. In simultaneous overexpression of the Jak-Stat target Ovo, tumors are larger and contain more cells per cyst. Scale bars, 10 µm **(H)** Quantification of cysts containing more than 64 cells, from **(G)**. Error bars, standard error. Scale bars, 10 µm.

Simultaneous overexpression of Dpp and Upd resulted in tumors containing many pMad^+^ cells (**Fig 4E**) and Stat^+^ cells (**Fig 4F**). Moreover, simultaneous expression of Bmp ligand Dpp together with EGT *ovo*, which is responsive to Jak-Stat but not Bmp signaling, resulted in enhancement of Bmp-induced tumor phenotype. Whereas Bmp tumors are typically comprised of spermatogonial-like cells with 32-64 germ cells per cyst, as previously shown^46^ (**Fig 4G**), simultaneous overexpression of Ovo and Dpp led to a larger testis with an increase in the number of germ cells found in each cyst (**Fig 4G-H**). Thus, addition of a Jak-Stat target, or Jak-Stat signaling itself, enhances the Bmp tumor phenotype rather than acting redundantly with the Bmp pathway. Together, these data suggest that Bmp and Jak-Stat signaling synergistically define germ cells’ behavior.

### Bmp-high, Jak-Stat low state is associated with dedifferentiation

The results thus far suggest that the EGTome is regulated in a modular manner by Bmp and Jak-Stat pathways, potentially permitting different outcomes based on reception of both, neither, or only one ligand. Combinations of signaling modules may enable equipotent cells (i.e. GSCs and spermatogonia) to choose between self-renewal, differentiation, or dedifferentiation. We tested this hypothesis by inducing mass dedifferentiation by transient ectopic expression of the differentiation factor Bam^23,24^ (*nos-gal4*, *tubulin*-*gal80, UAS-bam*, see method) (**Supp Fig 5A-C)**. We found that spermatogonia in mass dedifferentiation conditions that have not made attachment to hub cells had a high pMad signal, whereas they were not yet positive for Stat (**Figure 5A, Supp Fig 5D**). Stat signal returned in GSCs that were directly hub-proximal (**Figure 5A, Supp Fig 6A**). In control conditions in which spermatogonia are not massively dedifferentiating, spermatogonia were negative for both pMad and Stat. Together, this suggests that GSCs are defined by pMad^+^ and Stat^+^ (double positive), differentiating cells are defined by pMad^-^ Stat^-^ (dual negative), and the “third state” of dedifferentiation is associated with single-positive pMad^+^ Stat^-^.

**Figure 5.**
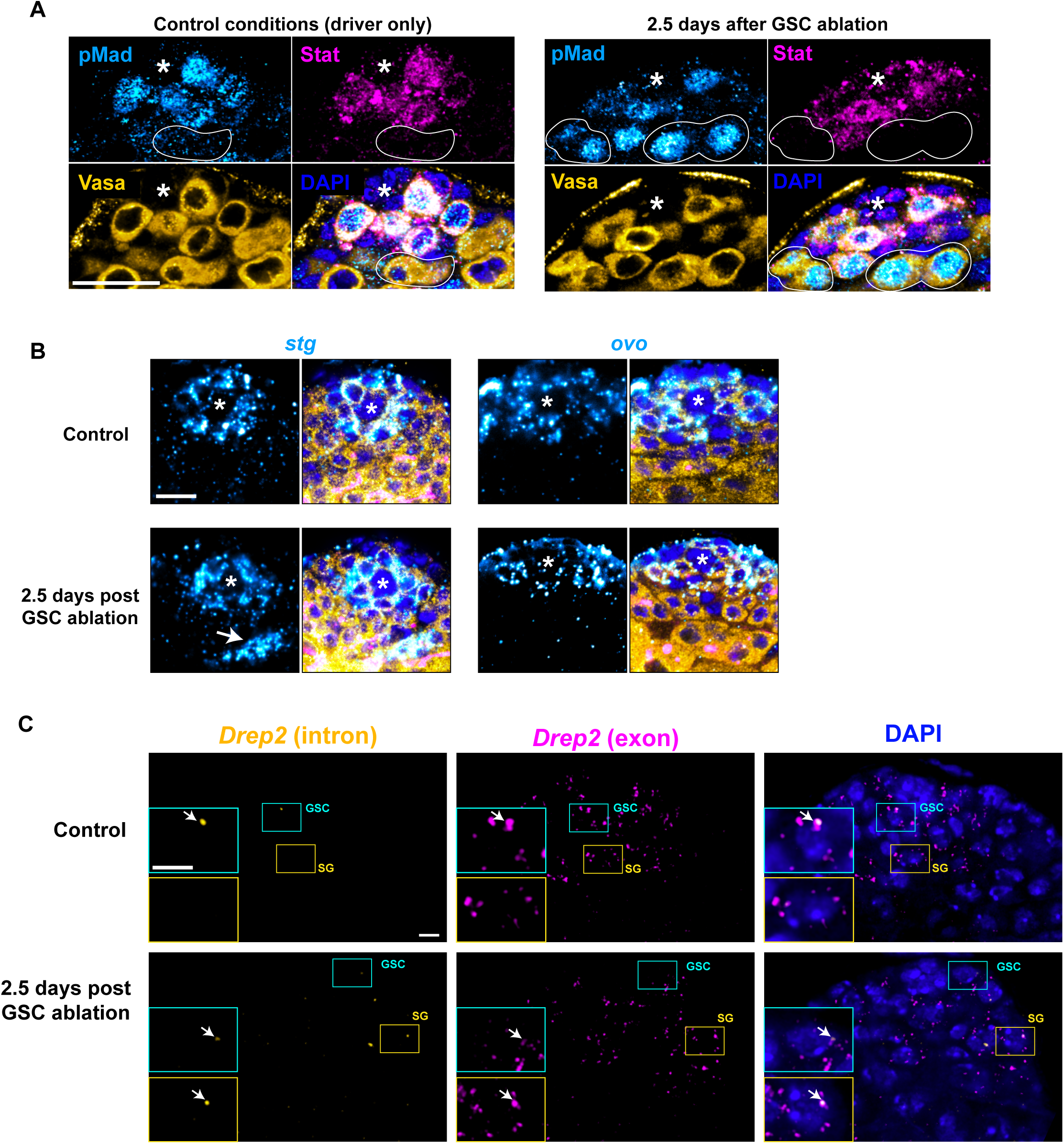
Dedifferentiation is characterized by a Bmp-on, Jak-Stat-off state in spermatogonia. **(A)** In control conditions, spermatogonia (outlined) are neither pMad^+^ or Stat^+^. 2.5 days after near-total GSC ablation via transient overexpression of Bam, dedifferentiation is frequent (see **Supp Fig 5A-C**). At this timepoint, spermatogonia (outlined) are pMad^+^, but not Stat^+^. Scale bars, 10 µm. **(B)** Bmp-sensitive transcript *stg* is found in later spermatogonia (arrow) in dedifferentiation conditions, but Bmp-insensitive transcript *ovo* is not. Scale bars, 10 µm. **(C)** Nuclear intronic signal of Bmp-sensitive EGT *Drep2* is restricted to GSCs in control conditions, but is present in spermatogonia in dedifferentiation conditions. Scale bars, 5 µm.

If Bmp reception accompanies the dedifferentiation state, increased presence of EGTs downstream of Bmp should be associated with dedifferentiation. Indeed, expression of Bmp-sensitive transcript *stg* (**Supp Fig 2A**, **Fig 3A**) was upregulated in later spermatogonia in dedifferentiation conditions as compared to control conditions (**Fig 5B**). By contrast, expression of *ovo*, which is a Jak-Stat-regulated EGT (**Supp Fig 2A**, **Fig 3C**), was not changed in dedifferentiation conditions (**Fig 5B**). Furthermore, active transcription of Bmp-sensitive EGT *Drep2* (**Figure 3B**), while typically restricted to GSCs (**Figure 2D**), was found in spermatogonia in dedifferentiation conditions (**Figure 5C**). These results suggest that the Bmp pathway activates dedifferentiation by promoting new Mad-dependent transcription of its downstream EGTs in spermatogonia.

These data are consistent with a previous observation that depletion of the RNA-binding protein Me31B resulted in mass dedifferentiation^39^ associated with upregulation of Mad – but not Stat^39^. Recapitulating the results from dedifferentiation arising from GSC loss (**Fig 5B**), we found that *me31b* knockdown likewise led to upregulation of *stg* (Bmp responsive EGT) but not *ovo (*Stat responsive EGT) in spermatogonia, suggesting that *me31b* RNAi recapitulates natural dedifferentiation with the activation of Bmp signaling and resulting transcript expression to promote spermatogonial dedifferentiation (**Supp Fig 6B**). Together, these data suggest that Bmp signaling and the resulting transcriptional output is a hallmark of spermatogonial dedifferentiation in both conditions of GSC loss and genetic perturbation.

Consistent with the notion that “Bmp on, Stat off” triggers dedifferentiation in spermatogonia that have not yet contacted the niche, the combination of Stat-off (*nos*>*stat^RNAi^*), Bmp-on (*nos*>*dpp*) resulted in high levels of dedifferentiation (**Fig 6A**). Dedifferentiation can be identified by the presence of hub-proximal spermatogonial cysts of ≥3 germ cells connected by fragmenting fusomes^24,39^. By contrast, differentiating spermatogonia typically have branched, continuous fusomes that connect all cells of the cyst^50^. Notably, the high proportion of dedifferentiation observed under Stat-off, Bmp-on condition (*nos*>*stat^RNAi^, dpp*) was likely not due to the loss of GSCs alone. Although low Jak-Stat signaling alone triggered partial GSC loss as reported previously^24^ (**Supp Fig 6C**), spermatogonial cysts attached to the hub under these conditions contained a branched fusome, indicative of differentiation, instead of dedifferentiation (**Supp Fig 5C-F**). This suggests that increased dedifferentiation found in Bmp-high, Jak-Stat-low conditions was not merely because of GSC loss from *stat^RNAi^*, but instead due to dedifferentiation triggered by Bmp overexpression. Although it was suggested that Jak-Stat activity promotes dedifferentiation^23,24^, the fact that Stat was never observed in dedifferentiating SGs that have not made contact with the hub may suggest that Stat is required to reestablish GSC identity upon returning to the niche, rather than initiating dedifferentiation. Consistent with this notion, in Bmp-high, Jak-Stat-low testes, Bmp-sensitive transcripts were expressed outside their normal GSC/early SG expression boundary; Bmp-insensitive transcripts, by contrast, retained their normal expression pattern (**Supp Fig 6G**). However, overexpression of individual Bmp-sensitive EGTs *Mov10* and *stg* was not sufficient to drive dedifferentiation, and the loss of Bmp-sensitive *Mov10* did not inhibit dedifferentiation (**Supp Fig 6 H-1**). Thus, the entire repertoire of Bmp-dependent transcription is likely required for dedifferentiation control, with individual EGTs insufficient in isolation. Together, these data suggest that a Bmp-high, Jak-Stat-low environment is sufficient to trigger dedifferentiation.

**Figure 6.**
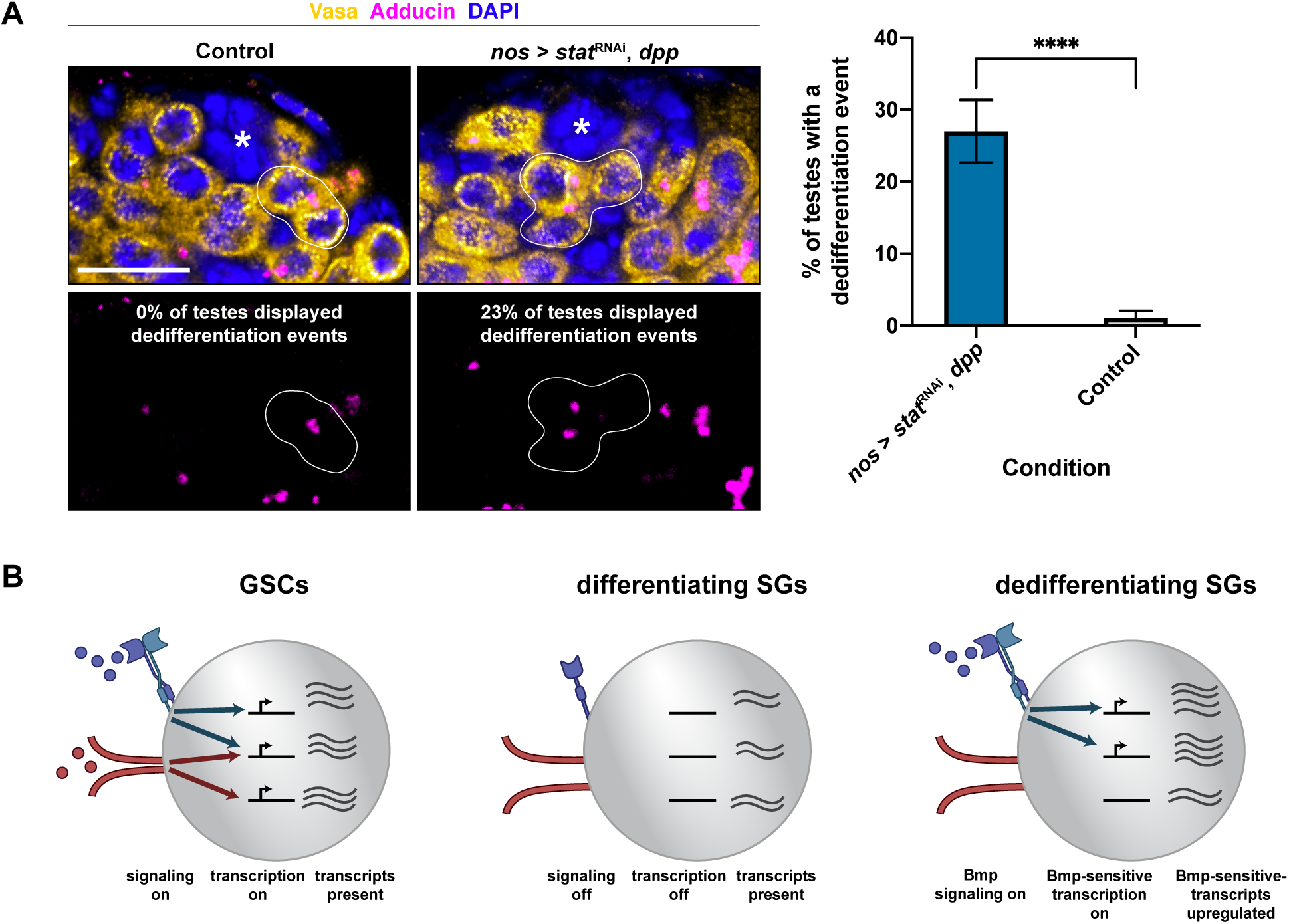
**(A)** Dedifferentiation is characterized by cysts of >2 germ cells with disconnected fusomes contacting the hub. This was observed frequently after simultaneous overexpression of Bmp ligand and RNAi of the Stat transcriptional effector. Outline indicates a dedifferentiating cyst. Scale bars, 10 µm. Error bars, standard error. ****, p<0.00001, unpaired t-test. **(B)** Model. GSC identity is set by the reception of niche signaling ligands, which trigger transcription of early germline transcripts and subsequent mRNA presence. During differentiation, these transcripts perdure and eventually dilute. However, there is no niche signaling reception, leading to a loss of a GSC-like chromatin state and a resulting lack of active EGT transcription. During dedifferentiation, Bmp, but not Jak-Stat ligand, is received, leading to new active transcription of Bmp-sensitive transcripts and third unique active transcriptional state.

## Discussion

The plasticity of adult stem cell populations, and the ability of partially differentiated cells to re-acquire stem cell character, has been increasingly recognized in diverse adult stem cell lineages^1^. In such a dynamic system, stem cells and differentiating cells *en route* to terminal differentiation must balance competing demands. Stem cells must remain distinct from their daughter cells, with a unique self-renewal identity and associated cell biological behaviors that support self-renewal. Differentiating cells must maintain stem cell potential even while adopting a differentiation trajectory. Our work revealed two parallel mechanisms to separate potential and trajectory (**Fig 6B**). First, our data suggest that stem cells are the only germline cells that actively transcribe the early germline transcriptome, whereas differentiating cells share the same set of transcripts as stem cells by inheriting them but not actively transcribing them. This may provide differentiating cells with the potential to revert back to stem cell identity without causing tumorous overproliferation. Second, the independent activity of the Bmp and Jak-Stat pathways allows GSCs and spermatogonia to balance three trajectories (i.e., self-renewal, differentiation, and dedifferentiation). Thus, we propose that transcriptome perdurance maintains potential in all GSCs and spermatogonia, whereas the dual niche signaling enables equipotent cells to make distinct choices (self-renewal vs. differentiation vs. dedifferentiation).

Notably, binary on/off activity of one factor alone cannot generate more than two states. However, combinatorial activity of two separate pathways generates a system in which reception of one, neither, or both each yields a distinct outcome. Prior work^48,49^ has suggested that the Stat and Bmp pathways depend on the activity of each other through cross-talk. In such a case, only two states could be defined by a dual signaling system. Our work suggests that the presence of two distinct and non-dependent pathways is essential for a stem cell niche that precisely balances self-renewal, dedifferentiation, and dedifferentiation. Correspondent with this model, we find that the Stat and Bmp pathways act independently and do not largely activate each other. Importantly, our determination of a GSC transcriptional “signature” – as opposed to a single marker – permitted the discovery that GSC identity is comprised of discrete transcriptional modules that require separable activators. Future work is likely to determine if the transcription factors of the Bmp and Jak-Stat pathways directly bind to all their EGT targets. Alternatively, Mad and Stat may bind to a subset of EGTs to initiate a transcriptional cascade, as some EGTs themselves encode for transcription factors.

Importantly, our study demonstrates both the benefits and limitations of using single-cell sequencing to categorize cell types. While such sequencing was vital to define the EGTome, without which it would be impossible to determine if Bmp and Jak-Stat signaling have distinct sets of targets, it also failed to distinguish GSCs and spermatogonia. Incomplete or incorrect inferences of cell type from transcriptomic studies are often attributed to translational regulation^51^. It awaits future investigation to test if EGTs are differentially translated or undergo different post-translational control in GSCs and spermatogonia. Indeed, restriction of pMad and Stat to GSCs likely occurs post-translationally^43,44^, and recent work has identified that translational control serves as a key regulator to distinguish somatic cyst stem cells and their progeny^52^. Our study in GSCs, however, suggests that ‘active transcription’, controlled by transcription factor activity, is an important and separable parameter of cells’ state. Whereas GSCs and early spermatogonia share a transcriptome (EGTome), GSCs are likely the only cell type that is actively producing EGTome, revealing an important limitation of single-cell transcriptomic analysis in isolation. Further analysis of GSCs vs. spermatogonia may reveal additional chromatin-level differences that reflect distinct active transcription without distinct transcriptomes.

Previously, we showed that GSCs are likely the only cells that can receive Bmp ligand secreted by the hub under normal conditions^53^. GSCs extend membranous protrusions (called microtubule-based nanotubes, or MT-nanotubes) into the hub cells, and the interface between the MT-nanotubes and the hub cell invaginations serves as the only site for productive ligand-receptor interaction. Therefore, spermatogonia displaced from the hub are excluded from engaging in the niche signaling^53^. The very presence or absence of GSCs could therefore determine the ability of spermatogonia to receive Bmp ligand. The Jak-Stat ligand Upd, by contrast, is known to be diffusion-limited through its association with the extracellular matrix^54^, and is likely never receivable by spermatogonia distal from the hub, regardless of presence or absence of GSCs. This allows for a precisely controlled emergent phenomenon, in which spermatogonia in the presence of GSCs do not receive a high Bmp signal and therefore do not activate a dedifferentiation program. In the absence of GSCs, Bmp ligand may diffuse and reach spermatogonia, initiating an active transcriptional program associated with dedifferentiation initiation. Jak-Stat ligand Upd is not received by these distal cells because it is diffusion-limited. Upon homing to the niche (i.e. reestablishment of attachment to the hub), such dedifferentiating spermatogonia will reactivate Jak-Stat signaling, thus returning to the true GSC state (by reacquiring GSC behavior and/or suppressing dedifferentiation-specific behavior such as migration).

A recent study^55^ proposed that Bmp ligand diffusion during homeostasis suppresses, rather than promotes, differentiation of the spermatogonia. Specifically, trapping Bmp ligand to the hub cell surface increased, rather than decreased, the frequency of spermatogonia dedifferentiating into GSC identity. Because Bmp ligand is known to exert distinct effects depending on its concentration^56^, trapping ligand on secreting cells may create a state of Bmp signaling in the spermatogonia distinct from direct perturbations of pathway components. It awaits future investigation to understand how specific levels of Bmp signaling, potentially in combination with varying levels of Jak-Stat signaling, may encode specific cellular states.

The reactivation of Bmp targets during dedifferentiation has another important implication. Dedifferentiation in an unperturbed tissue to replace sporadic GSC loss is likely supported by dedifferentiation of one- or two-cell spermatogonia, and GSCs may be homeostatically maintained through interconversion with their immediate daughters^5,45^. However, after GSC loss, dedifferentiation of 4-cell spermatogonial cysts is frequently seen^24^. These one- or two-cell spermatogonia differ from 4-cell spermatogonia in terms of number of divisions from being a GSC and in distance from the hub. Our work identifies another difference in these populations: high expression of the EGTome. Because our GSC-enriched transcripts are constantly seen in early spermatogonia (**Fig 1** and **Supp Fig 1**), inheritance of EGTome from GSCs may maintain a pool of particularly dedifferentiation-competent 2-cell cysts. By contrast, Bmp-sensitive GSC-enriched transcripts are found after GSC loss in 4-cell spermatogonial cysts (**Fig 5**). Therefore, this *de novo* transcription of Bmp-responsive factors could be particularly critical during mass dedifferentiation after GSC loss, whereas high perdurant transcript levels primarily maintain dedifferentiation competence during homeostasis.

Taken together, this work provides important insights into the puzzle of how stem cell lineages balance self-renewing and differentiating cell populations, while maintaining the opportunity for interconversion between these populations. Maintaining this balance, such that dedifferentiation does not occur too frequently in homeostasis resulting in tumorigenesis, yet can be robustly triggered after GSC loss, is paramount to tissue equilibrium.

## Materials and Methods

### Fly husbandry and strains

Unless otherwise stated, flies were raised at 25° and assayed at age 1-4 days. All flies were raised on standard Bloomington media without propionic acid. For a list of all fly strains used in this study, see **Table S2.** To generate dedifferentiation conditions by overexpression of Bam in GSCs, we took advantage of the temperature-sensitive Gal80 system to drive Bam. Specifically, *nos-gal4*, *tubulin-gal80* flies were crossed with *UAS-bam* flies and raised at 18°. Upon eclosion, adult *nos-gal4, tubulin-gal80, UAS-bam* flies and their control siblings were transferred to a 29° water bath for 64 hours. This timepoint was empirically determined to provide maximum GSC loss (see Supp Fig 5) without overwhelming spermatogonial loss. Following the 29° incubation, flies were transferred back to 18° for an additional 64 hours before dissection. This time was likewise determined to allow the maximum number of confident dedifferentiation events.

### Identification of EGTs

Because of the low cell count in the final clustering, differential gene expression analysis was performed using the SCDE package that is most reliable for low cell counts^30^. First, an expression matrix was created with “positive” cells (putative GSCs) and “negative” cells (putative spermatogonia) from cell names (colnames) within the cds object. A csv file was generated from the merged feature-barcode matrix in bash the mat2csv command (CellRanger). SCDE was then run as per package defaults. The output file was merged with the original features file to annotate gene short names and FlyBase ID. The hits were sorted by “conservative estimate” (ce). See Table S1 for the full list. See the Supplementary information for the detailed protocol for scRNA sequencing.

### Microscopy and image processing

All microscopy images were taken on a Leica Stellaris using a 63x oil-immersion objective and processed using FIJI. For the 3D reconstruction, the Imaris machine-learning based 3D modeling was used. The detailed protocols for immunofluorescence staining and FISH are provided in the Supplementary information.

## Data and materials availability

Replicates 1-5 of the single-cell sequencing data are available at GEO under accession number GSE311622. Replicate 6 of the single-cell sequencing data was previously published^28^ and is available at GEO under accession number GSE220615.

## Acknowledgments

We acknowledge members of the Yamashita lab and Erika Bach for critical input. We thank FlyBase, Bloomington Drosophila Stock Center, and the Drosophila Genomics Resource Center for reagents. YMY was supported in this work by the Howard Hughes Medical Institute and the Gordon and Betty Moore Foundation. AAR was supported in this work by 1F32GM143850-01 and 1K99GM154143-01A1.

## Author Contributions

Conceptualization, AAR and YMY; investigation, AAR and HH; analysis, AAR; writing – original draft, AAR; writing – editing, AAR and YMY; funding acquisition, AAR and YMY; supervision, YMY.

