## Supplemental Materials for "Niche-dependent modular regulation of the stem cell transcriptome separates cell identity and potential"

### Supplemental Figures and Tables, Materials and Methods

### Supplementary Figures

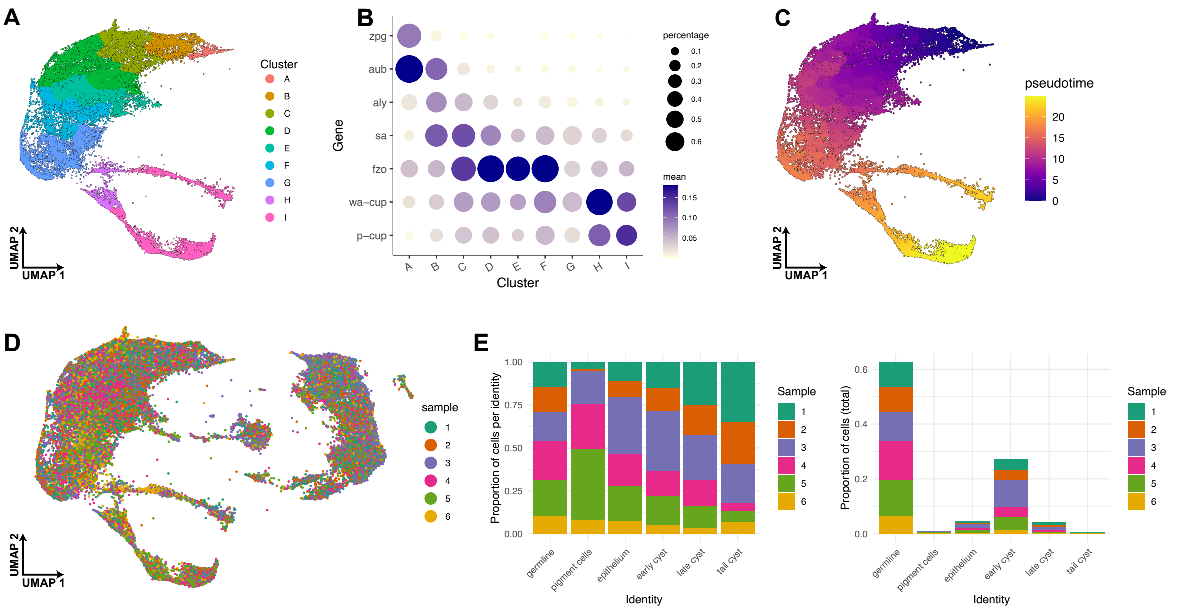


**Supplemental Figure 1. Characterization of single-cell sequencing data and validated EGTs. (A)** UMAP representation of all germline annotated cells in the testis. Color indicates merged and quality-controlled germline cluster annotation, as in **Fig 1C**. Clusters labeled alphabetically. **(B)** Dotplot displaying clusters, as in **(A)**, with expression of known differentiation-stage markers indicated. Size of dot indicates the percentage of cells in that cluster with an expression level above 0 for that gene. Color indicates the mean expression value across all cells in that cluster for that gene. Transcript presence correlates with known differentiation stages. **(C)** Calculated pseudotime values of germline cells. Germline cells form an unbroken trajectory. Cluster A was imputed as the root. **(D)** UMAP representation of all testis cells. Cells colored by sample from the 6 biological replicates. **(E)** Proportion of cells from each cluster from each biological replicate. Each replicate is represented across all clusters, and outliers are largely only seen in very low-number clusters (pigment cells and tail cyst cells, especially).

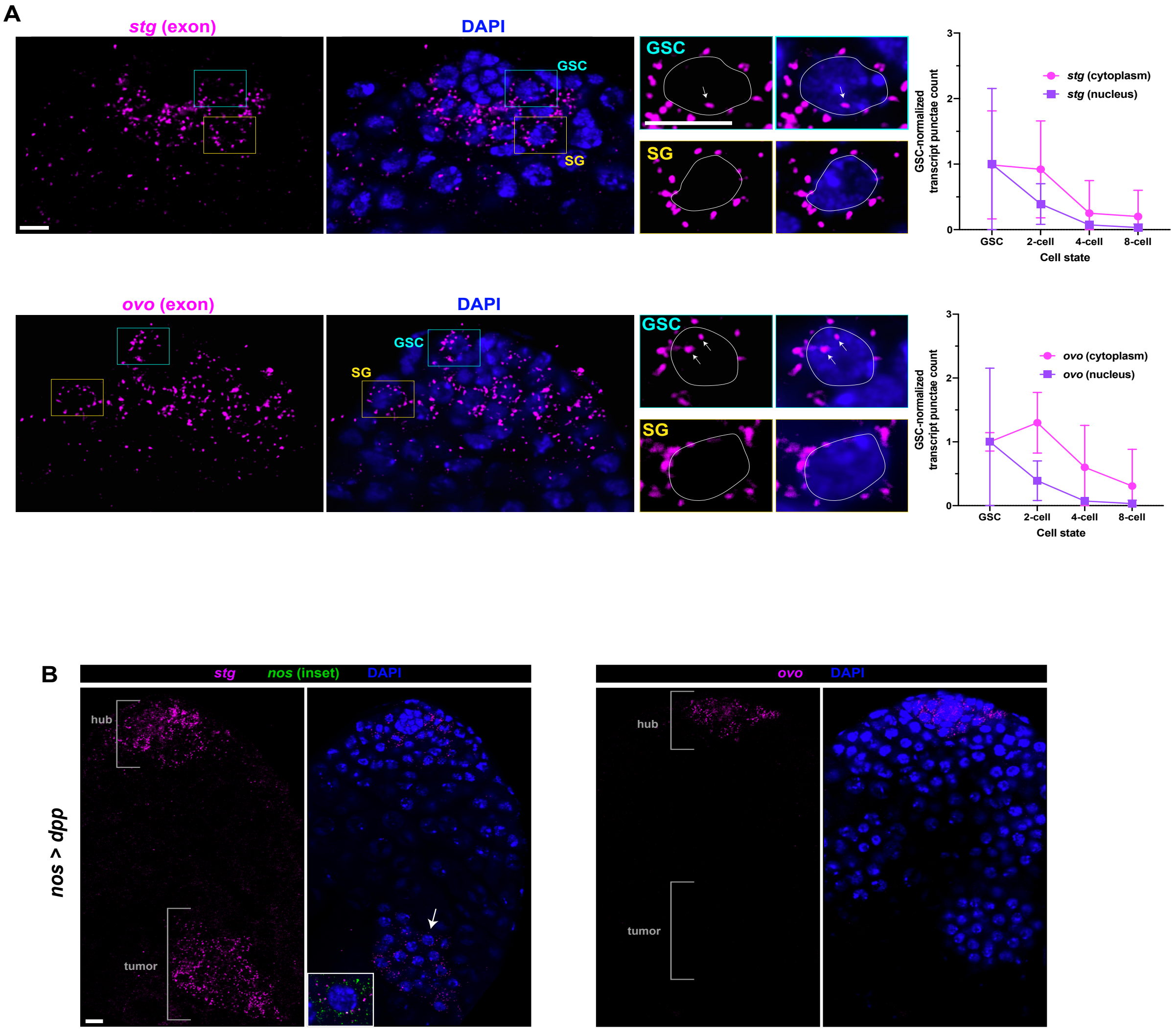


**Supplemental Figure 2.Characterization of EGT *stg* and *ovo* expression by single-molecule FISH. . (A)** Nascent transcription can be inferred by nuclear exonic signal. Nuclear signal of EGTs *stg* and *ovo* is largely restricted to GSCs. Scale bars, 5 μm. Error bars, standard error. **(B)** Single-molecule FISH demonstrates that *stg* is expressed in hub-distal Bmp tumor cells, but *ovo* is only expressed in hub-proximal cells in its normal expression pattern. Tumor expression of *stg* is in *nanos*$^{+}$ germ cells (inset). Scale bars, 10 μm.

A

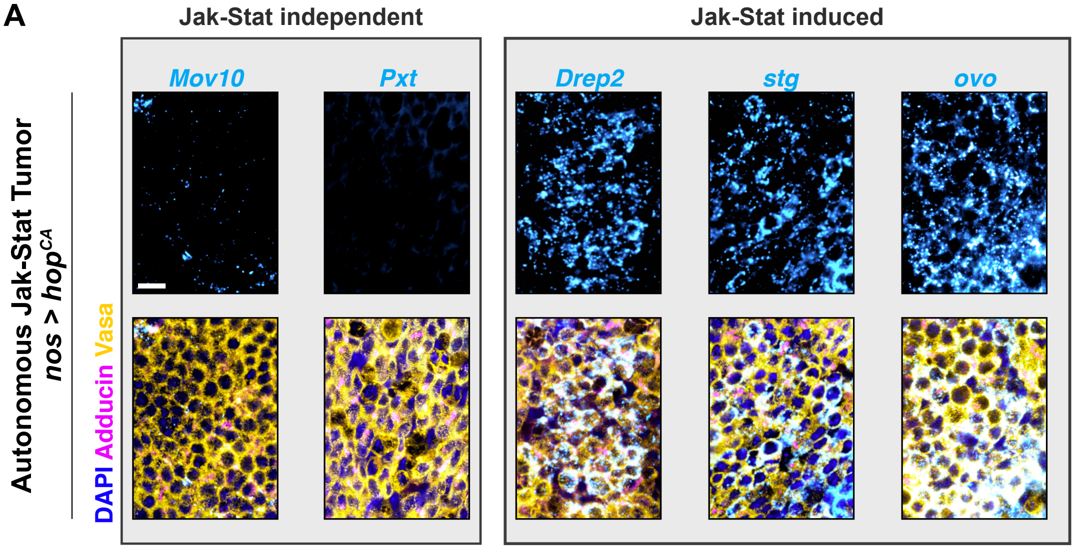
Jak-Stat independent
Jak-Stat induced
Mov10
Pxt
Drep2
stg
ovo
Autonomous Jak-Stat Tumor
nos > hopCA
DAPI Adducin Vasa

B

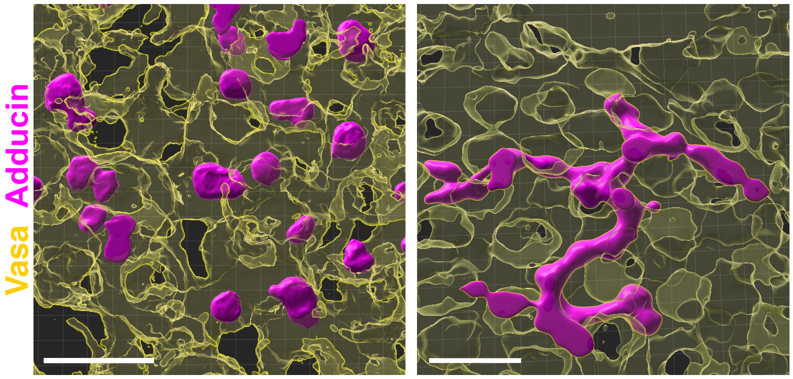
Vasa Adducin

C

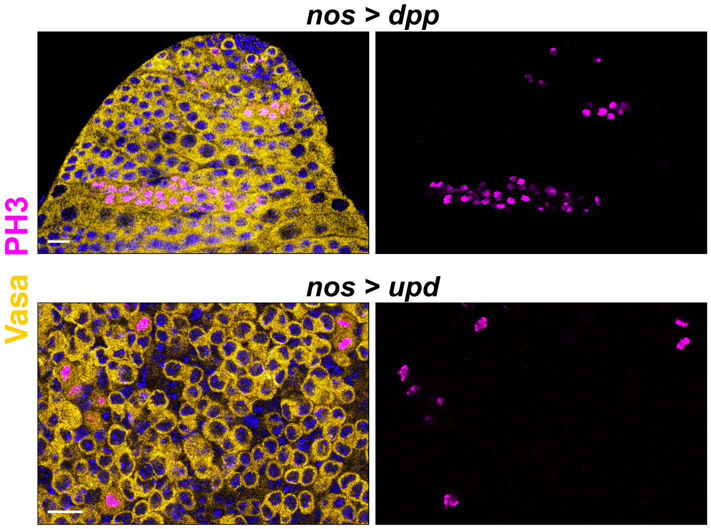
nos > dpp
nos > upd
PH3
Vasa

**Supplemental Figure 3. Bmp and Jak-Stat tumors differ. (A)** Tumors are generated by germ cell autonomous expression of a constitutively active form of the Jak Kinase (Hopscotch). EGTs *Mov10* and *Pxt* are not largely present in these germ cell tumors, whereas EGTs *Drep2*, *stg*, and *ovo* are. This mirrors their expression in tumors that arise from overexpression of Upd. Scale bar, 10μm. **(B)** 3D reconstruction (see methods) of a Jak-Stat tumor (left) and a Bmp tumor (right). In the Jak-Stat tumor, fusomes (Adducin$^{+}$ structures) are individual and small. In the Bmp tumor, fusomes have a large branched morphology. **(C)** Mitotic cells are marked by phosphorylated histone 3. In Bmp tumors, cells in a cyst enter meiosis synchronously. In Jak-Stat tumors, cells enter meiosis asynchronously with their neighbors.
Scale bars, 10 μm.

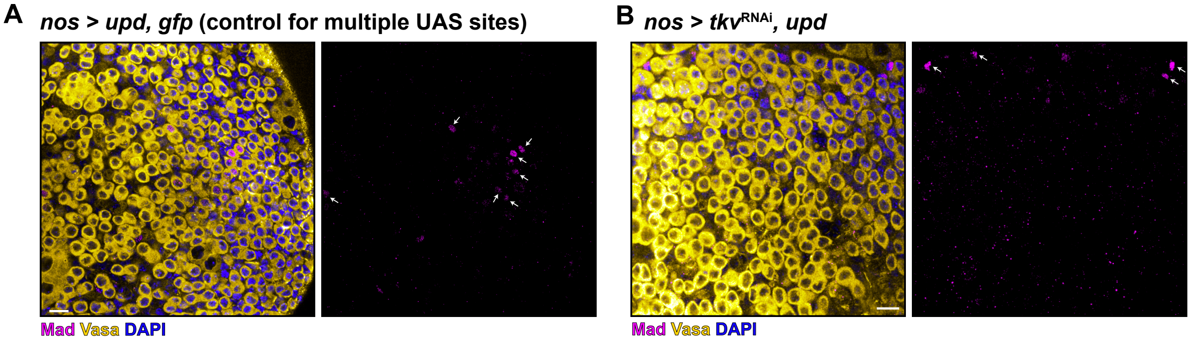


**Supplemental Figure 4. Formation of Jak-Stat tumors does not require Bmp signaling. (A)** In Jak-Stat tumors, sparse pMad$^+$ germ cells are seen (arrows), especially around hub-like structures. GFP was simultaneously driven with Upd to control for dilution of the Gal4 by multiple UAS sites. **(B)** When Upd overexpression is combined with RNAi of the Dpp receptor Thickveins, tumors of GSC-like cells still form. Now, pMad is confined to cyst cells (arrows) and is not found in germ cells.

Scale bars, 10 µm.

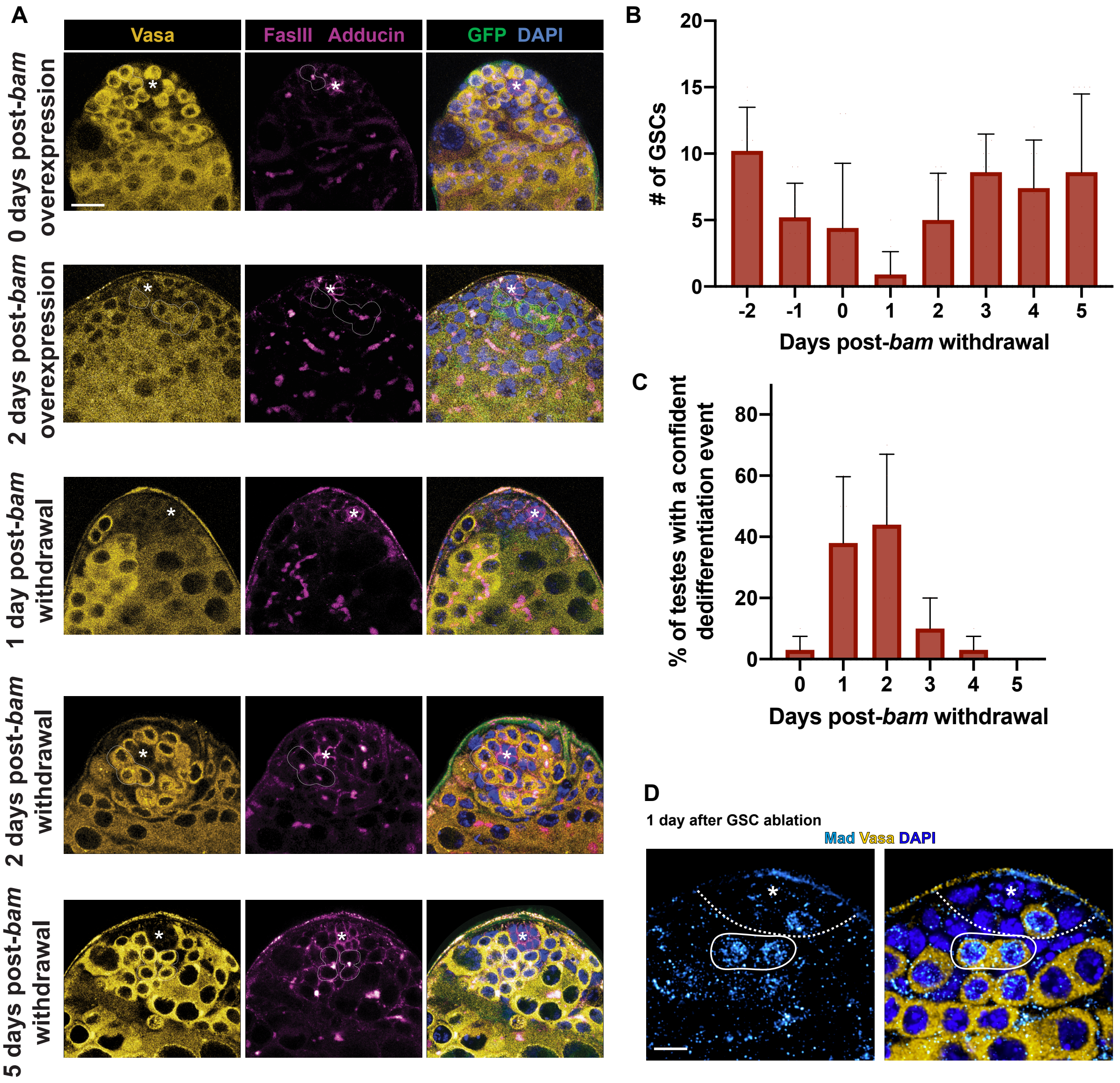


**Supplemental Figure 5. Dedifferentiation timecourse by ectopic Bam expression. (A)** Top row: Prior to ectopic bam expression, germ cells (Vasa+) surround the hub (FasIII+, asterisk). GSCs directly contact the hub. GSC divisions (dotted line) that generate a differentiating germ cell are transiently connected by a fusome (Adducin+ focus). Second row: After forced bam expression, few GSCs remain. Ectopic bam expression indicated with GFP (green). Third row: All GSCs are gone soon following withdrawal of ectopic bam. Fourth row: After withdrawal of ectopic bam expression, spermatogonia will begin dedifferentiating, as indicated by connected spermatogonia in which one cell has contacted the niche and is connected to >1 other cells by discontinuous fusomes. Bottom row: Five days after withdrawal of ectopic bam, the niche has been repopulated by GSCs, and these dedifferentiated GSCs have begun dividing to produce more differentiating daughter cells. Scale bar, 10 µm **(B)** Quantification of GSC number after withdrawal of ectopic Bam. GSCs have depleted by day 1 post-withdrawal, and have returned to normal levels by day 3. Error bar, standard error. **(C)** Quantification of "confident dedifferentiation events," as defined by ≥3 germ cells connected by fragmented fusomes contacting the hub. Likely other dedifferentiation events exist that have not yet contacted the hub, and/or have not yet fragmented the fusome. Such confident dedifferentiation events peak at approximately 2 days after withdrawal of ectopic Bam. **(D)** Example of pMad in spermatogonia in an early stage of dedifferentiation prior to GSCs largely repopulating the niche. One GSC lies inside the dotted line, proximal to the hub (asterisk). Spermatogonia distal to the hub (circle) are positive for pMad.

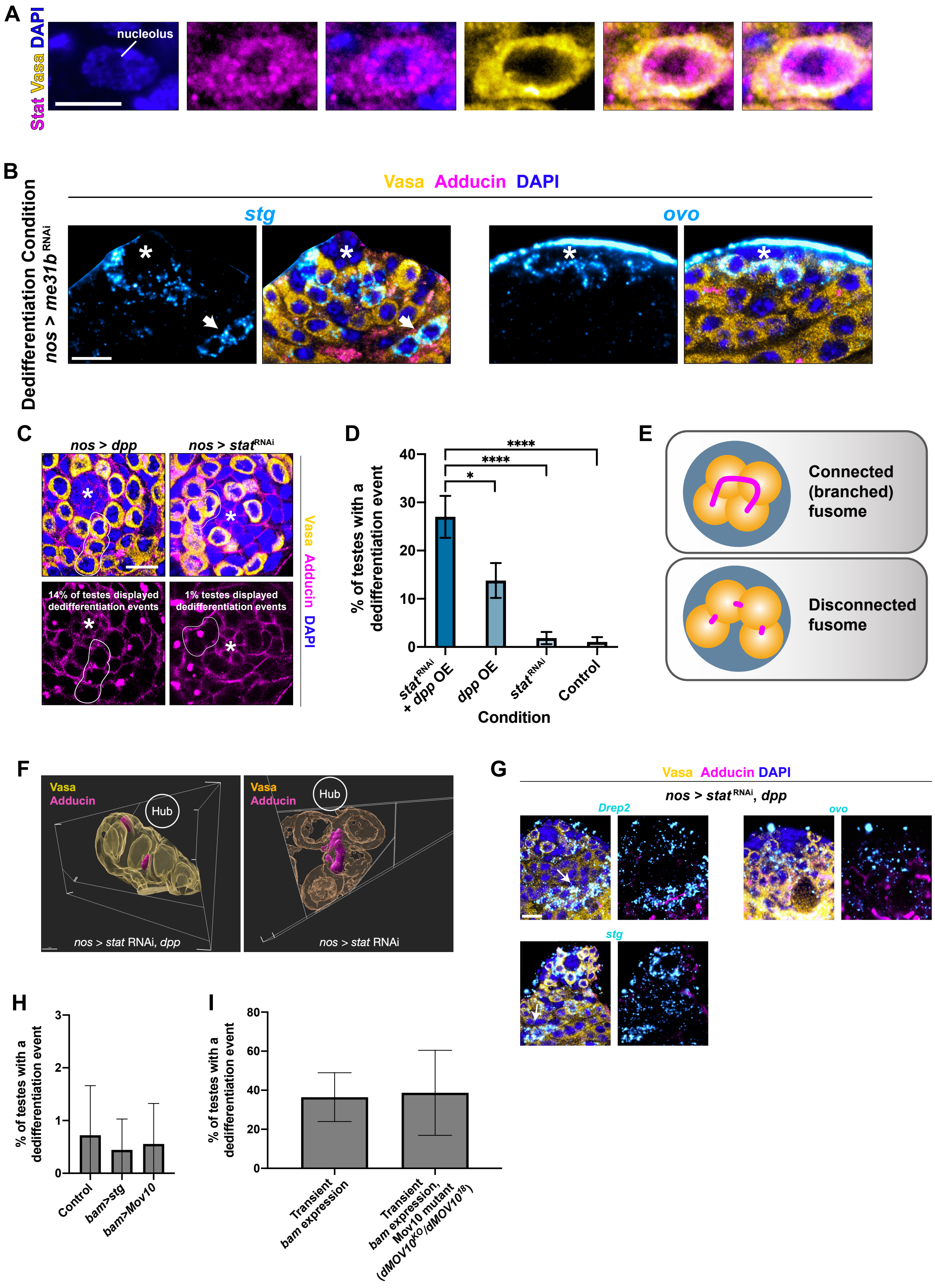


**Supplemental Figure 6. Characterization of dedifferentiation conditions. A)** Inset from **Fig 5A**. Stat signal in GSCs is both nuclear and cytoplasmic, but excluded from the nucleolus. Vasa signal is entirely cytoplasmic. Scale bar, 5 µm **(B)** Dedifferentiation can be triggered by RNAi of *me31b*. In this condition, Bmp-sensitive transcript *stg* appears in spermatogonia outside its normal expression zone, but Bmp-insensitive transcript *ovo* does not. Scale bar, 10 µm **(C)** Dpp tumor testes have an increased number of somatic hub cells and an increased number of GSCs[46]. In this condition, we observed 14% of testes with dedifferentiation events, as identified by ≥3 germ cells connected by fragmented fusomes contacting the hub (circled). Knockdown of *stat*, by contrast, leads to a reduced number of GSCs in the hub. However, we did not observe considerable dedifferentiation events above control, only asymmetric division events (circled). Scale bar, 10 µm **(D)** Quantification of data in **Supp Fig 5D** and **Fig 5C**. First and last columns are also present in **Fig 5C**. Error bars, standard error. * p<0.01, **** p<0.00001, unpaired t-test. **(E)** Cartoon representation of spermatogonia with connected (branch) fusome and disconnected fusome. **(F)** 3D reconstruction of germ cells in $stat^{RNAi}$ and $stat^{RNAi}$, *dpp* conditions. In $stat^{RNAi}$, *dpp* testes, germ cells cysts adjacent to the hub had disconnected fusomes connecting ≥3 germ cells. In $stat^{RNAi}$ alone conditions, however, germ cell cysts contacting the hub had connected fusomes. This may suggest that Jak-Stat signaling plays a role in a return to GSC identity once the process of dedifferentiation has returned dedifferentiating spermatogonia to the hub. **(G)** *Drep2* and *stg* are both Bmp-sensitive transcripts and display expression outside of hub-adjacent GSCs in $stat^{RNAi}$, *dpp* conditions. *ovo*, by contrast, is Bmp-independent, and does not show such outside expression. *ovo* expression is also reduced in GSCs in this condition, suggesting some dependence on Jak-Stat signaling. **(H-I)** Quantification of dedifferentiation events after perturbation of individual Bmp-sensitive EGTs. EGTs *stg* and *Mov10* do not individually affect dedifferentiation rates in basal conditions (**H**) or in mass dedifferentiation conditions (**I)**

| Table S1: Single Cell Differential Expression (SCDE) analysis of GSC-enriched factors | | | | | | | |
|---|---|---|---|---|---|---|---|
| **FB ID** | **Gene short name** | **lower bound** | **maximum liklihood estimate** | **upper bound** | **conservative estimate** | **uncorrected Z score** | **corrected Z score** |
| **FBgn0003525** | **stg** | **-7.4024** | **-6.5583** | **-5.4220** | **-5.4220** | **-5.4543** | **-4.6328** |
| FBgn0001120 | gnu | -7.1427 | -6.2986 | -5.3570 | -5.3570 | -6.0586 | -5.2427 |
| **FBgn0003028** | **ovo** | **-6.8830** | **-6.1687** | **-5.1622** | **-5.1622** | **-5.1550** | **-4.3125** |
| FBgn0261574 | kug | -6.6232 | -5.9414 | -4.9999 | -4.9999 | -5.1868 | -4.3469 |
| FBgn0004167 | kst | -6.4934 | -5.8440 | -4.9350 | -4.9350 | -5.5590 | -4.7365 |
| FBgn0010382 | CycE | -6.4934 | -5.8116 | -4.7401 | -4.7401 | -4.9717 | -4.1244 |
| FBgn0020248 | stet | -6.3960 | -5.7466 | -4.6752 | -4.6752 | -4.7558 | -3.9088 |
| FBgn0015371 | chn | -6.6882 | -5.9090 | -4.6427 | -4.6427 | -5.2199 | -4.3786 |
| **FBgn0261987** | **Pxt** | **-6.2012** | **-5.4869** | **-4.5129** | **-4.5129** | **-4.8592** | **-4.0104** |
| FBgn0003118 | pnt | -7.0453 | -6.1038 | -4.5129 | -4.5129 | -5.0345 | -4.1877 |
| FBgn0052088 | sunn | -6.2986 | -5.6492 | -4.5129 | -4.5129 | -5.0045 | -4.1587 |
| FBgn0033402 | Myd88 | -7.3050 | -6.4284 | -4.5129 | -4.5129 | -4.4523 | -3.5913 |
| FBgn0011659 | Mlh1 | -6.2012 | -5.5194 | -4.3181 | -4.3181 | -3.6729 | -2.7768 |
| **FBgn0028408** | **Drep2** | **-9.0258** | **-7.6946** | **-4.2856** | **-4.2856** | **-3.9691** | **-3.0831** |
| **FBgn0034187** | **Mov10** | **-6.5258** | **-5.7466** | **-4.2531** | **-4.2531** | **-4.8251** | **-3.9766** |
| FBgn0085434 | NaCP60E | -6.3310 | -5.6492 | -4.1882 | -4.1882 | -4.2327 | -3.3607 |
| FBgn0085312 | CG34283 | -6.0388 | -5.2272 | -4.1233 | -4.1233 | -4.8496 | -4.0006 |
| FBgn0027339 | jim | -6.8505 | -6.0064 | -4.1233 | -4.1233 | -3.9535 | -3.0673 |
| FBgn0003023 | otu | -6.2012 | -5.4544 | -4.0259 | -4.0259 | -4.5899 | -3.7347 |
| FBgn0250819 | CG33521 | -6.2012 | -5.3895 | -3.9934 | -3.9934 | -4.1227 | -3.2453 |
| FBgn0039883 | RhoGAP100F | -6.2012 | -5.4544 | -3.9285 | -3.9285 | -3.3805 | -2.4706 |
| FBgn0038273 | CG14860 | -5.8765 | -5.2921 | -3.8960 | -3.8960 | -4.0437 | -3.1600 |
| **FBgn0014141** | **cher** | **-6.0713** | **-5.3570** | **-3.8311** | **-3.8311** | **-3.7028** | **-2.8061** |
| FBgn0264491 | how | -8.2141 | -7.2726 | -3.7661 | -3.7661 | -3.8109 | -2.9157 |
| **FBgn0001981** | **esg** | **-5.9090** | **-5.2596** | **-3.7337** | **-3.7337** | **-3.3918** | **-2.4835** |
| FBgn0036433 | CG9628 | -8.0193 | -5.0648 | -3.7337 | -3.7337 | -3.7000 | -2.8037 |
| FBgn0003984 | vn | -6.7531 | -5.2272 | -3.7012 | -3.7012 | -3.6873 | -2.7918 |
| FBgn0032026 | CG7627 | -6.0713 | -5.3895 | -3.6687 | -3.6687 | -3.6085 | -2.7052 |
| FBgn0033785 | Sans | -6.5258 | -5.7142 | -3.6687 | -3.6687 | -3.7633 | -2.8678 |
| FBgn0004569 | aos | -7.1102 | -5.3246 | -3.6038 | -3.6038 | -2.8685 | -1.9103 |
| FBgn0002592 | E(spl)m2-BFM | -7.6622 | -5.0648 | -3.5713 | -3.5713 | -3.0229 | -2.0817 |
| FBgn0023214 | edl | -7.1102 | -5.2272 | -3.5389 | -3.5389 | -3.5299 | -2.6235 |
| FBgn0010453 | Wnt4 | -7.3375 | -4.9350 | -3.5064 | -3.5064 | -3.6201 | -2.7184 |
| FBgn0035049 | Mmp1 | -6.5583 | -5.1298 | -3.5064 | -3.5064 | -3.4923 | -2.5832 |
| FBgn0259876 | Cap-G | -6.6232 | -5.9414 | -3.4739 | -3.4739 | -4.9492 | -4.1001 |
| FBgn0052814 | CG32814 | -6.3310 | -5.5518 | -3.4739 | -3.4739 | -3.3428 | -2.4324 |
| FBgn0037999 | CG4860 | -6.2661 | -5.6168 | -3.4739 | -3.4739 | -3.5663 | -2.6600 |
| FBgn0030960 | Atg101 | -6.0713 | -5.4220 | -3.4415 | -3.4415 | -3.4359 | -2.5261 |
| FBgn0266418 | wake | -6.1362 | -5.3895 | -3.4415 | -3.4415 | -3.2496 | -2.3321 |
| FBgn0267033 | mamo | -8.7985 | -8.2466 | -3.4090 | -3.4090 | -7.0524 | -6.1703 |

| | | | | | | | |
|---|---|---|---|---|---|---|---|
| FBgn0263392 | Tet | -7.6946 | -4.4479 | -3.4090 | -3.4090 | -3.2265 | -2.3067 |
| FBgn0000562 | egl | -6.1038 | -5.1298 | -3.4090 | -3.4090 | -2.5487 | -1.5826 |
| FBgn0029648 | CG3603 | -5.8765 | -5.3246 | -3.4090 | -3.4090 | -3.1720 | -2.2462 |
| FBgn0023023 | CRMP | -6.0388 | -5.3895 | -3.3765 | -3.3765 | -3.4337 | -2.5240 |
| FBgn0263077 | CG43340 | -6.4284 | -5.3895 | -3.3765 | -3.3765 | -3.3441 | -2.4333 |
| FBgn0261244 | inaE | -6.6232 | -5.8765 | -3.3116 | -3.3116 | -3.9257 | -3.0379 |
| FBgn0037215 | beta-Man | -6.2012 | -5.5194 | -3.2791 | -3.2791 | -3.4294 | -2.5192 |
| FBgn0004872 | piwi | -7.6622 | -6.9479 | -3.2142 | -3.2142 | -4.6592 | -3.8089 |
| FBgn0265974 | ttv | -6.5583 | -5.8765 | -3.2142 | -3.2142 | -4.7798 | -3.9300 |
| FBgn0285925 | Fas1 | -8.1816 | -7.3050 | -3.1493 | -3.1493 | -4.9015 | -4.0519 |
| FBgn0010434 | cora | -6.3960 | -5.2272 | -3.1493 | -3.1493 | -3.4121 | -2.5044 |
| FBgn0013763 | Idgf6 | -6.4934 | -5.0973 | -3.0519 | -3.0519 | -2.8682 | -1.9103 |
| FBgn0039167 | CG17786 | -5.9414 | -5.2272 | -3.0194 | -3.0194 | -3.2692 | -2.3519 |
| FBgn0030808 | RhoGAP15B | -6.1038 | -5.2921 | -2.9869 | -2.9869 | -3.0795 | -2.1438 |
| FBgn0262737 | mub | -8.0518 | -7.3050 | -2.9869 | -2.9869 | -6.4688 | -5.6148 |
| FBgn0001085 | fz | -5.9739 | -5.2596 | -2.9545 | -2.9545 | -3.0852 | -2.1501 |
| FBgn0267912 | CanA-14F | -7.0128 | -6.2336 | -2.9220 | -2.9220 | -4.6619 | -3.8114 |
| FBgn0086698 | frtz | -5.9739 | -5.2921 | -2.9220 | -2.9220 | -2.9652 | -2.0169 |
| FBgn0260400 | elav | -5.9414 | -5.2921 | -2.8895 | -2.8895 | -2.8750 | -1.9168 |
| FBgn0051373 | CG31373 | -6.9479 | -6.0064 | -2.8571 | -2.8571 | -4.0957 | -3.2170 |
| FBgn0038418 | pad | -5.9090 | -5.0324 | -2.8246 | -2.8246 | -3.0848 | -2.1500 |
| FBgn0002121 | l(2)gl | -7.1752 | -6.3635 | -2.7597 | -2.7597 | -4.2735 | -3.4019 |
| FBgn0031294 | IA-2 | -7.1102 | -6.3310 | -2.7597 | -2.7597 | -5.1338 | -4.2904 |
| FBgn0015396 | jumu | -7.4024 | -6.6882 | -2.7272 | -2.7272 | -5.4599 | -4.6372 |
| FBgn0029849 | Efr | -5.7791 | -5.0324 | -2.7272 | -2.7272 | -2.9616 | -2.0142 |
| FBgn0003416 | sl | -5.7466 | -5.0648 | -2.6947 | -2.6947 | -2.6605 | -1.6915 |
| FBgn0035812 | CG7457 | -6.5258 | -5.8440 | -2.6947 | -2.6947 | -4.5749 | -3.7197 |
| FBgn0034501 | CG13868 | -5.9414 | -4.8375 | -2.6947 | -2.6947 | -2.3545 | -1.3877 |
| FBgn0030941 | wgn | -6.0713 | -5.3570 | -2.6623 | -2.6623 | -2.9055 | -1.9531 |
| FBgn0029830 | Grip | -6.2336 | -5.4869 | -2.6298 | -2.6298 | -2.8591 | -1.9009 |
| FBgn0040602 | CG14545 | -7.7920 | -3.9934 | -2.5973 | -2.5973 | -4.6251 | -3.7740 |
| FBgn0041604 | dlp | -6.2012 | -5.3895 | -2.5973 | -2.5973 | -3.4616 | -2.5535 |
| FBgn0038045 | NANS | -5.8440 | -4.8700 | -2.5649 | -2.5649 | -2.7114 | -1.7428 |
| FBgn0026263 | bip1 | -6.2012 | -5.4869 | -2.4999 | -2.4999 | -3.2563 | -2.3395 |
| FBgn0011236 | ken | -7.2076 | -6.4609 | -2.4999 | -2.4999 | -4.5780 | -3.7219 |
| FBgn0266521 | stai | -4.6103 | -3.5064 | -2.4675 | -2.4675 | -5.8205 | -4.9893 |
| FBgn0278608 | Dsp1 | -3.8635 | -3.1168 | -2.4025 | -2.4025 | -7.1606 | -6.2058 |
| FBgn0033649 | pyr | -6.0388 | -5.1622 | -2.4025 | -2.4025 | -2.9486 | -2.0004 |
| FBgn0001104 | Galphai | -4.4479 | -3.3765 | -2.3701 | -2.3701 | -5.7317 | -4.8978 |
| FBgn0037703 | JHDM2 | -5.8765 | -5.2272 | -2.3376 | -2.3376 | -2.8613 | -1.9022 |
| FBgn0022893 | Df31 | -3.5713 | -2.8571 | -2.3051 | -2.3051 | -7.1608 | -6.2058 |
| FBgn0037779 | CG12811 | -6.5258 | -5.7466 | -2.3051 | -2.3051 | -3.8120 | -2.9165 |
| FBgn0086909 | CG31751 | -6.0064 | -5.2596 | -2.2727 | -2.2727 | -3.0703 | -2.1341 |
| FBgn0027585 | CG8740 | -5.9090 | -5.0973 | -2.2402 | -2.2402 | -2.9154 | -1.9649 |

| | | | | | | | |
|---|---|---|---|---|---|---|---|
| FBgn0262656 | Myc | -3.6687 | -2.9545 | -2.2402 | -2.2402 | -7.1608 | -6.2058 |
| FBgn0000635 | Fas2 | -4.0908 | -3.1168 | -2.2402 | -2.2402 | -6.6926 | -5.8365 |
| FBgn0029976 | snz | -6.0064 | -5.3570 | -2.2402 | -2.2402 | -2.8609 | -1.9022 |
| FBgn0052251 | Claspin | -7.5648 | -6.7856 | -2.2077 | -2.2077 | -4.0357 | -3.1519 |
| FBgn0035770 | pst | -6.2336 | -5.4869 | -2.1753 | -2.1753 | -3.2874 | -2.3703 |
| FBgn0032587 | CG5953 | -7.9868 | -3.6363 | -2.1428 | -2.1428 | -4.3542 | -3.4876 |
| FBgn0011704 | RnrS | -7.7596 | -7.0453 | -2.1428 | -2.1428 | -5.2057 | -4.3661 |
| FBgn0031540 | Pif1 | -6.4609 | -5.6817 | -2.1103 | -2.1103 | -4.0228 | -3.1378 |
| FBgn0026143 | CDC45L | -6.1038 | -5.3246 | -2.1103 | -2.1103 | -3.1716 | -2.2462 |
| FBgn0285926 | Imp | -3.1817 | -2.5973 | -2.0454 | -2.0454 | -7.1608 | -6.2058 |
| FBgn0001624 | dlg1 | -7.4024 | -3.2791 | -2.0454 | -2.0454 | -5.0260 | -4.1794 |
| FBgn0038476 | kuk | -3.7661 | -2.9545 | -2.0454 | -2.0454 | -6.2051 | -5.3794 |
| FBgn0037531 | CG10445 | -6.6232 | -5.8440 | -2.0454 | -2.0454 | -4.3000 | -3.4303 |
| FBgn0024980 | Syx4 | -6.1038 | -5.3246 | -2.0454 | -2.0454 | -3.3626 | -2.4511 |
| FBgn0265523 | Smr | -3.7337 | -2.7272 | -2.0129 | -2.0129 | -7.1608 | -6.2058 |
| FBgn0030520 | Pdcd4 | -3.1493 | -2.4999 | -2.0129 | -2.0129 | -7.1608 | -6.2058 |
| FBgn0038197 | foxo | -7.5323 | -6.7531 | -2.0129 | -2.0129 | -3.3815 | -2.4715 |
| FBgn0035626 | lin-28 | -7.0453 | -5.2596 | -2.0129 | -2.0129 | -3.2314 | -2.3122 |

| Table S2: Resource Table | | | |
|---|---|---|---|
| **Type** | **Resource** | **Source** | **Use** |
| Drosophila strain | nos-gal4 | PMID: 9501989 | Early germ cell driver |
| Drosophila strain | UAS-dpp | BDSC 1486 | Bmp overexpression |
| Drosophila strain | UAS-upd | PMID: 10346822 | Jak-Stat overexpression |
| Drosophila strain | UAS-OvoB | BDSC 38430 | Ovo overexpression |
| Drosophila strain | UAS-Stat RNAi | BDSC 33637 | RNAi of *Stat* |
| Drosophila strain | USA-me31b RNAI | BDSC 38923 | RNAi of *me31b* |
| Drosophila strain | UAS-tkv RNAi | BDSC 40937 | RNAi of *tkv* (Bmp receptor) |
| Drosophila strain | UAS-HopCA | Ref #59 | Germ cell autonomous Jak-Stat overexpression |
| Drosophila strain | UAS-stg | BDSC 4777 | Stg overexpression |
| Drosophila strain | UAS-Mov10 | Ref #33 | Mov10 overexpression |
| Drosophila strains | dMov10K0, dMov1018 | Ref #33 | Mov10 mutant |
| Antibody | Rat anti-vasa | DSHB | Immunofluorescent labeling of germ cells. Used 1:20 |
| Antibody | Rabbit anti-Mad | ab52903 | Immunofluorescence. Used 1:200 |
| Antibody | Guinea Pig anti-Stat | PMID: 26131929 | Immunofluorescence. Used 1:200 |
| Antibody | Mouse anti-Adducin | DSHB | Immunofluorescent labeling of the fusome. Used 1:20 |
| Antibody | Rabbi anti-PH3 | Sigma (06-570) | Immunofluorescent labeling of mitotic cells. Used 1:200 |
| Plasmid | esg plasmid | DGRC-FI20009 | Probe generation |
| Plasmid | Drep2 plasmid | DGRC-LD32009 | Probe generation |
| Plasmid | Pxt plasmid | DGRC-LD43174 | Probe generation |

| | | | |
|---|---|---|---|
| Plasmid | cher plasmid | DGRC-RE44980 | Probe generation |
| Plasmid | stg plasmid | DGRC-LD47579 | Probe generation |
| Plasmid | ovo plasmid | DGRC-LD47350 | Probe generation |
| Plasmid | Mov10 plasmid | DGRC-LD34829 | Probe generation |

### Supplementary Materials and Methods

#### scRNA-seq

Tissue isolation and dissociation for scRNA-seq. Dissociation was performed as previously described[28]. Briefly, on the day of the dissection, fresh dissociation buffer (2 mg/mL collagenase in Trypsin LE) was prepared. Testes from 1-4 day old flies were dissected in 1x phosphate-buffered saline (PBS) and immediately transferred to cold PBS tubes for a maximum of 30 minutes while the remaining dissections were performed. Testis samples were centrifuged at 135 rcf, PBS was removed and replaced with dissociation buffer. Testes were incubated in maceration buffer for 30 minutes with gentle vortexing every 10 minutes at room temperature. Following incubation, samples were pipetted up and down until no visible tissue chunks remained. Sample was then passed through a 35 µm filter and centrifuged at 135 rcf for 7 minutes. The supernatant was removed and the pellet was resuspended in 1 mL of calcium/magnesium-free Hanks' Balanced Salt Solution (HBSS). The sample was centrifuged for a final time at 135 rcf for 7 minutes and all but 50 µL of the HBSS was removed. The pellet was resuspended in the remaining 50 µL. Cell viability and density was assayed on a hemocytometer with DIC imaging and Acridine Orange stain.

Library preparation and single-cell sequencing. Cells were processed using the 10X Genomics Chromium Controller and Chromium Single Cell Library And Gel Bead Kit following standard manufacturer's protocol. Amplified cDNA libraries were quantified by bioanalyzer and size-selected using AMPure beads. Samples were sequenced on a NovaSeq S4

Mapping, dimensionality reduction, clustering, and pseudotime analysis. Reads were mapped to the DM6 reference genome using the 10X CellRanger pipeline with default parameters, on a per-sample basis. The resulting feature matrices for all 6 biological samples were read into the R package Monocle3[29] using load_cellranger_data. The samples were combined using combine_cds and aligned using align_cds; no other sample integration methods were used. The data was preprocessed using preprocess_cds and dimensionality reduction was performed with reduce_dimensions. Initial partitioning was performed by clustering cells at resolution $1^{-5}$. At this point, the germline partition was embedded and a principal graph was generated using learn_graph. The germline partition was then further subclustered with resolution 0.0006 using cluster_cells again. These clusters were then manually merged and recoded, as in Supp Fig 1A. The pseudotime trajectory was inferred using order_cells of the germline partition, with "cluster A" identified as the root based on expression of known early germline markers, including *zpg, nanos,* and *aub*. Finally, cluster A was isolated and subclustered one last time at resolution 0.01. Additional subclustering at higher resolutions did not reproducibly generate similar clusters when different random seeds were set; therefore, this was considered the last possible biologically relevant clustering for this dataset.

**Tissue staining for imaging**

Immunofluorescence. Immunofluorscence staining was performed as described previously[27]. Briefly, testes were dissected in PBS, transferred to 4% formalydehyde in PBS and fixed for 30 minutes. Tissues were then washed in PBS + 0.1% Triton X (PBST) for 3 times, 10' each, followed by incubation with primary antibody in 3% BSA in PBST at 4° overnight. Samples were washed in again 3x10' in PBST and incubated in secondary antibody in 3% BSA in PBST. Washes were performed again and samples were mounted in Vectashield with DAPI. The exception to this protocol was for the anti-pMad antibody. For stains with this antibody, two modifications were made. First, initial washes were performed with PBS + 0.3% Triton X prior to incubation in primary. Second, a 30' blocking step in 5% BSA (not 3%) was performed, and the primary antibody incubation was in 5% BSA. When immunofluorescence was combined with TSA FISH, the FISH protocol was performed first, followed by the immunofluorescence protocol. The pMad antibody recognizes phosphorylated (active) Mad specifically, whereas the Stat antibody recognizes both active and inactive Stat. Stat presence in the nucleus and cytoplasm is demonstrated in Supp Fig 6.

TSA FISH. All RNA *in situ* experiments, with the exception of Fig 1G, Supp Fig 2A, and Fig 2C, were performed using tyramide signal amplification (TSA) FISH. RNA oligoprobes were synthesized from Drosophila Genome Resource Center (DGRC) plasmids, as described[57] (**Table S2**). Briefly, plasmids were linearized and amplified as follows: pBS backbone with SK-30 (F) and SKMet (R), pFLC-1 packbone with M13(-21) (F) and M13 (REV) (R), and pOT2a backbone with PM001a (F) and PM002a (R). *in vitro* transcription reactions were performed on the PCR products with the following polymerases: T3 for the pBS or pFLC-1 backbones, and SP6 for the pOT2a backbone. The reactions were performed with Digoxigenin (DIG)-conjugated ribonucleotides (Roche). Following the transcription reaction (overnight at 37°), the reaction was treated with DNase and then probes were precipitated with 7.5M Ammonium Acetate and 100% ethanol, and resuspended in formamide.

FISH with tyramide signal amplification was then performed as described[28]. Testes were isolated and fixed in 4% formaldehyde for 30–60 min, dehydrated into methanol and stored at –20 °C for up to 1 month. After rehydration in PBS +0.1% Triton- X100, testes were permeabilized with 4 μg/ml Proteinase K for 6 min and washed with Pre- Hybridization solution (50% deionized formamide, 5x SSC, 1 mg/ml yeast RNA, 1% Tween- 20) for up to 2 hr at 56°. Testes were incubated overnight at 56 °C with probes diluted 1:800 in Hybridization solution (50% deionized formamide, 5x SSC, 1 mg/mL yeast RNA, 1% Tween- 20, and 5% Dextran Sulfate). After washes in Pre- Hybridization solution, 2 x SSC, and 0.2 x SSC, then PBS +0.1% Triton- X100, samples were blocked for 30 min in 1% Roche Western Blocking Reagent prior to incubating overnight at 4 °C with either anti- FITC with horseradish peroxidase conjugate (Roche) at a 1:2000 concentration or anti- DIG- with horseradish peroxidase conjugate (Roche) at a 1:1500 concentration in 1% Roche Western Blocking Reagent. Fluorescent tyramide development and amplification were performed by first placing the testes for 5 min in borate buffer (0.1 M boric acid, 2 M NaCl, pH 8.5), followed by 10 min in borate buffer with rhodamine (1:1000) or fluorescein (1:1500) tyramide, and 0.0003% hydrogen peroxide. Samples were washed before continuing to immunofluorescence.

Stellaris FISH. For single-molecule FISH (Fig 1G and Supp Fig 2A), FISH with probes from Biosearch Technologies was performed as previously described[58]. Probes were designed against the coding sequences of *nanos*, *ovo*, *Mov10*, and *stg* using the Biosearch Technologies design tool. Testes were dissected in 1x PBS and fixed in 4% formaldehyde in 1x PBS for 30 min. Then testes were washed briefly in 1x PBS and permeabilized in 70% ethanol overnight at 4°. Testes were briefly rinsed with wash buffer (2x SSC, 10% formamide) and then hybridized overnight at 37° in hybridization buffer containing 2x SSC, 10% dextran sulfate (Sigma-Aldrich), 1mg/ml Escherichia coli tRNA (Sigma-Aldrich), 2 mM Vanadyl Ribonucleoside complex (NEB), 0.5% BSA (Ambion), and 10% formamide. Fluorescently labeled probes were added to the hybridization buffer to a final concentration of 100 nM. Following hybridization, samples were washed three times in wash buffer for 20 min each at 37°.

HCR FISH. For the nascent transcript FISH on *Drep2* (Fig 2D and 5C), we used Hybridization Chain Reaction (HCR) FISH. Using the Molecular Instruments design tool, nascent probes were designed against the third and fourth introns and exonic probes were designed against the coding sequence of *Drep2*. Testes were dissected in 1x PBS and fixed in 4% formaldehyde in 1x PBS for 20 minutes and then washed twice, 5 minutes per wash, in 1x PBS 0.1% Tween-20. Permeabilization was achieved by washing for 2 hours in 1x PBS + 0.1% Triton X-100. The testes were then washed twice, 5 minutes per wash, in 5x SSC + 0.1% Tween-20. Probes were added to Hybridization Buffer to a final concentration of 10nM. Samples were hybridized overnight at 37C shaking. Samples were washed four times, 15 minutes per wash, at 37° with Probe Wash Buffer (Molecular Instruments). During the second wash, hairpin solutions for desired amplifiers (Molecular Instruments) were created by individually heating each hairpin at 95C for 90 seconds in a PCR machine and allowing the samples to slowly cool to room temperature. Amplifiers were added to Amplification Buffer (Molecular Instruments) to a final concentration of 60nM. Samples were washed twice, 5 minutes per wash, with 5x SSC 0.1% Tween-20, and then transferred to Amplification Buffer + amplifiers and incubated overnight at room temperature nutating. Testes were then washed twice, 30 minutes per wash, with 5x SSC 0.1% Tween-20.
